# A comprehensive benchmark of graph-based genetic variant genotyping algorithms on plant genomes for creating an accurate ensemble pipeline

**DOI:** 10.1101/2023.07.19.549631

**Authors:** Ze-Zhen Du, Jia-Bao He, Wen-Biao Jiao

**Affiliations:** National Key Laboratory for Germplasm Innovation and Utilization of Horticultural Crops, Huazhong Agricultural University, Wuhan, China; Hubei Hongshan Laboratory, Wuhan, China

**Keywords:** Genome graph, Plant genomes, Genotyping, Structural variation, Benchmarking

## Abstract

**Background:** Although sequencing technologies have boosted the measurement of the sequencing diversity of plant crops, it remains challenging to accurately genotype millions of genetic variants, especially structural variations, with only short reads. In recent years, many graph-based variation genotyping methods have been developed to address this issue and tested for human genomes, however, their performance in plant genomes remains largely elusive. Furthermore, pipelines integrating the advantages of current genotyping methods might be required, considering the different complexity of plant genomes.

**Results:** Here we comprehensively evaluate eight such genotypers in different scenarios in terms of variant type and size, sequencing parameters, genomic context, and complexity, as well as graph size, using both simulated and read data sets from representative plant genomes. Our evaluation reveals that there are still great challenges to applying existing methods to plants, such as excessive repeats and variants or high resource consumption. Therefore, we propose a pipeline called Ensemble Variant Genotyper (EVG) that can achieve better genotype concordances without increasing resource consumption. EVG can achieve comparably higher genotyping recall and precision even using 5× reads. Furthermore, we demonstrate that EVG is more robust with an increasing number of variants, especially for insertion and deletion.

**Conclusions:** Our study will provide new insights into the development and application of graph-based genotyping algorithms. We conclude that EVG provides an accurate, unbiased, and cost-effective way for genotyping both small and large variations and will be potentially used in population-scale genotyping for large, repetitive, and heterozygous plant genomes.

## Background

Genetic variants are typically divided into Single Nucleotide Polymorphism (SNP), insertion or deletion (indels, 1-49 bp), and Structural Variation (SV, ≥50 bp, including insertion, deletion, inversion, duplication, translocation, and complex rearrangements) based on their size and type [1, 2]. With the advances of high-throughput sequencing technologies, studies such as the 1000 Genomes Project and the Rice 3K Genomes Project have released large amounts of genetic variations, which contribute to the studies of pan-genomes, genome-wide associations, population genetics, and domestication [3–6]. One of the essential requirements for these studies is the rapid and correct genotyping (determination of genotypes) of millions of genetic variations for hundreds or thousands of individuals [3–6]. Conventional genotyping strategies usually rely on short-read mapping against a linear reference genome [2, 7–9]. However, these methods often introduce alignment errors due to reference bias, leading to erroneous genotypes for some variants, particularly those from regions highly divergent from the reference [10, 11]. In particular, genotyping SVs remains extremely challenging in population-scale studies where many individual genomes are sequenced solely using short-read sequencing technologies [3, 5, 11].

Recent advancements in pangenome graph-based genotyping algorithms are expected to mitigate reference bias and enhance the accuracy of genotyping across all types of genetic variations [12–15]. In such a graph, nodes typically represent the sequences, and edges indicate the connections between the sequences. Variations manifest as “bubbles”, and a path through the graph can be transformed into a haplotype sequence that represents combinations of different sequence variations [15]. By incorporating the reference genome as well as non-reference alleles into sequence or variation graphs, these algorithms can precisely genotype variations for individual genomes based on short-read data. They use either read (e.g., VG and GraphTyper2) or k-mer (e.g., BayesTyper and PanGenie) alignments against the graphs to achieve high accuracy [16–19]. However, the complex pangenome graph also expands the search space for read mapping. For instance, the original VG algorithm (VG-MAP) maps short reads to arbitrary variation graphs using generalized compressed suffix arrays to remove reference bias and improve alignment accuracy. Nevertheless, it is at least an order of magnitude slower than linear genome mappers, making it challenging to apply to large genomes or complex graphs [16]. A more recent version, VG-Giraffe, based on the seed-and-extend algorithm, can accelerate mapping [20]. Unlike VG-MAP and VG-Giraffe, which map reads to the whole-genome graph, only aligning reads to variant breakpoints (such as GraphTyper2) or comparing read k-mer coverages at k-mer represented variants (such as BayesTyper and PanGenie) can also reduce runtime [17–19]. Besides, the mapping accuracy of reads decreases as the number of nodes in the graph increases [21].

Remarkably, most of these algorithms were initially developed and tested for human genomes [16–18, 20, 22–24]. Although some graph-based genotypers such as VG have been applied to variation genotyping for crop genomes like rice [25], soybean [26], tomato [27], etc., the detailed performance of these tools remains elusive for plants as the complexity of plant genomes varies greatly in terms of genome size, repeat content, heterozygosity, and polyploidy. For example, repeat-enriched SVs can introduce inaccurate coverages of k-mers or reads at variant sites, thereby affecting the performance of graph-based genotyping methods that rely on such coverages for genotyping [18, 19, 23].

To address these issues, we first investigate the impact of read length and depth, number of variants, repeat density, heterozygous rate, etc. on existing graph-based genotypers in plant genomes [16–20, 24, 28]. Our findings suggest that there are still some challenges in applying existing methods to plants, such as excessive repeats and variants or high resource consumption. To overcome these challenges, we present an Ensemble Variant Graph-based tool, EVG, which can accurately genotype SNPs, indels, and SVs using short reads. Compared to other graph-based genotypers, EVG achieves higher genotyping accuracy and recall with only 5× sequencing data. Furthermore, the genotyping of EVG remains robust even as the number of nodes in the pangenome graph increases.

## Results

### Graph-based variant genotyper selection

To our knowledge, there are currently twelve graph-based genotyping tools available **(Table S1)**. For this study, we selected eight open-source graph-based genotyping tools that broadly fall into two categories: read alignment based (including VG-MAP [16], VG-Giraffe [20], Paragraph [24], GraphTyper2 [17] and Gramtools [29]) and k-mers alignment based (including BayesTyper [18] and PanGenie [19]) **(Table 1)**. We also conducted experiments to assess the performance of GraphAligner [28], a graph-based aligner that can utilize graphs from VG for alignment. Other tools were excluded from this study either because they are currently unsuitable for plant genomes (such as HISAT [30] and Minos [31]) or because they cannot genotype all types of genetic variations (like KAGE [32]). Among them, Minos is designed for bacterial genomes, while HISAT-genotype requires reconstruction of a typing database and complex conversion for plants in the algorithm.

**Table 1.** Overview of algorithms or models used in different graph-based genotyping tools.

| Software | Graph types | Indexing | Mapping | Genotyping | Reference |
| --- | --- | --- | --- | --- | --- |
| VG-MAP | Variation graph (bi-directed) | GCSA2 | Read alignment (Seed-cluster-chain, GSSW) | Coverage of local haplotypes | Hickey, G. et al. [16] |
| BayesTyper | Variation graph (DAG) | k-mer hash table | Read k-mer (n-best) | Generative model of the sequencing process with noise and diplotype k-mer counts | Sibbesen, J.A. et al. [18] |
| Paragraph | Variation graph (DAG) | k-mer hash table | Read alignment (S&E, GSSW) | Relative likelihood of local haplotypes based on coverages of breakpoints | Chen, S. et al. [24] |
| GraphTyper2 | Variation graph (DAG) | k-mer hash table | Read alignment (S&E) | Relative likelihood of local haplotypes based on breakpoint coverages and coverage in/de-crease | Eggertsson, H.P. et al. [17, 22] |
| VG-Giraffe | Variation graph (bi-directed) | GBWT, Minimizer | Read alignment (Seed-cluster-chain, GBWT) | Coverage of local haplotype | Siren, J. et al. [20] |
| Gramtools | Variation graph (nested DAG) | vBWT | Read alignment (S&E) | Likelihood of allele based on base-level and allele-level coverage | Letcher, B. et al. [29] |
| PanGenie | Variation graph (DAG) | k-mer hash table | Read k-mer | HMM of local haplotypes with k-mer counts and recombination rates | Ebler, J. et al. [19] |
| GraphAligner (+ VG) | Variation or overlap graph (bidirected) | Minimizer (default) | Read alignment (S&E) | VG used | Rautiainen, M. et al. [28] |

Note: DAG: directed acyclic graph, S&E: Seed-and-Extend, GCSA: Generalized Compressed Suffix Array, GBWT: Graph Burrows-Wheeler Transform, HMM: Hidden Markov Model, vBWT: variation-aware Burrows-Wheeler Transform.

These tools utilize different graph indexing approaches to improve alignment efficiency and/or support multiple graph manipulations **(Table S1)**. Specifically, VG employs GBWT [33], GCSA2 [34], and Minimizers [35] for graph storage and searching, whereas BayesTyper relies on k-mer-based graph indexing. To avoid potential memory overload, BayesTyper leverages Bloom filters to screen read k-mers, storing only those present in the haplotype [18]. Similarly, PanGenie adopts k-mer-based graph indexing using De Bruijn graphs. Subsequently, the software either aligns reads to nodes or directly counts k-mer coverage [19]. The genotyping results are then probabilistically scored based on statistical distribution modeling of observed and noise read/k-mer coverages.

### Overall performance on simulated data

To evaluate the performance of the tools, we first constructed a comprehensive simulation panel. Considering that plant genomes vary widely in size and repeats [36, 37], a series of data sets of paired-end shorted reads were simulated for each of four representative plant genomes (*Arabidopsis thaliana*, *Oryza sativa*, *Zea mays* and *Brassica napus*) with different genome sizes (135–2,300 Mb) and repeat contents (21.42-88.9%) **(Table S2)**. To generate simulated short reads from alternative (no-reference) genomes, we introduced different types (SNP, indels (<50 bp), and SVs (≥50 bp)) and numbers of variants into the reference genome of each plant species [5, 38–41] (see Methods for details). We repeat such a simulation of paired-end short reads with varying read lengths (100 bp, 150 bp, 250 bp), insert sizes (300 bp, 400 bp, 500 bp, 600 bp) and sequencing depths (5×, 10×, 20×, 30×, 50×) **(Table S3)**. Precision and recall were used to assess the genotyping performance of different software, and ROC curves were drawn according to the genotype quality or read depth.

As graph-based genotyping tools can leverage sequence information from multiple genomes, we first determined the overall performance of these tools based on the genome graph from eight *A. thaliana* genomes (one reference genome assembly, variants from other seven genomes, and 30× paired-end short reads) **(Fig. 1 a-c and Fig. S1a)**. For genomes to be genotyped, we simulated 467,512 SNPs, 38,207 indels, 4,572 insertions, 4,346 deletions, 232 inversions, and 100 duplications **(Table S4)**. Note that all these variants are incorporated into the genome graph. For SNP genotyping, all tools demonstrate high precision (>0.97), while only BayesTyper (0.99), Paragraph (0.98), and GraphTyper2 (0.93) have a recall rate greater than 0.90 **(Fig. S1a)**. Similarly, nearly all tools show high precision rates (0.80-0.99) and relatively lower recall rates (0.81-0.98) for indels genotyping. However, only BayesTyper (0.98), PanGenie (0.99), Gramtools (0.98), and Paragraph (0.97) maintain precision above 0.95 **(Fig. 1a)**. The performance of genotyping large insertions and deletions (≥50 bp) varies greatly among the tools. Despite different recall rates, all tools except Gramtools present genotyping precision above 0.8 **(Fig. 1b and 1c)**. Overall, Paragraph, GraphTyper2, and BayesTyper outperformed other tools in terms of genotyping performance **(Fig. 1b and 1c)**.

**Fig. 1.**
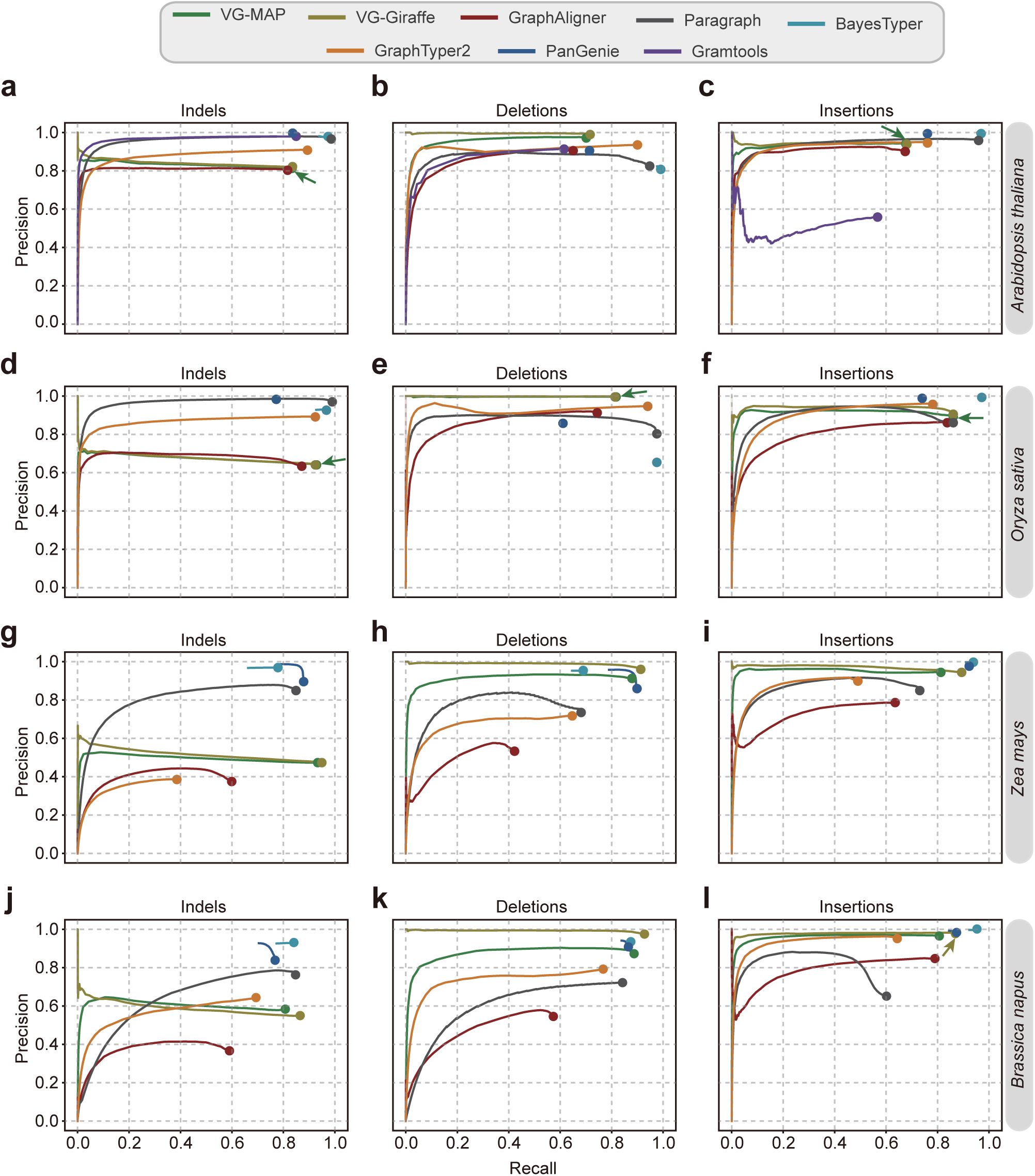
Overall genotyping performance of different graph-based tools based on simulated data. The genome graphs of *Arabidopsis thaliana* **(a, b, c)**, *Oryza sativa* **(d, e, f)**, *Zea mays*, **(g, h, i)** and *Brassica napus* **(j, k, l)**, are constructed based on one reference genome and seven alternative genomes derived by introducing known variants into the reference genome. Paired-end (2×150 bp) short reads with 30× depth are simulated for genotyping. For each genotyper, precision is plotted against recall as the genotyping quality threshold varies. Read depth (DP) on variant sites is used as a proxy score when genotype quality (GQ) is not available. Arrows indicate the circles hidden by other circles in the plot due to identical or nearly identical precision values.

We also evaluated these tools using the simulated data from rice, maize, and rapeseed genomes **(Tables S2 and S3)**. Note that Gramtools is excluded from the assessment for other plant genomes due to potential issues related to excessive chromosome length. We observed a similar recall of genotyping all types of variants in rice genomes as in *A. thaliana*, but with slightly decreased precision. **(Fig. 1d-f and Fig. S1c)**. For genotyping in the maize genome, VG-MAP and VG-Giraffe are the only tools that maintain high precision (0.91-1.00) and recall (0.82-0.95) in SV genotyping, while BayesTyper and PanGenie present high precision (0.97, 0.89) and recall (0.78, 0.88) for indel genotyping **(Fig. 1g-i and Fig. S1e)**. The genotyping performance in *Brassica napus* genomes was even lower, particularly for SNPs and indels **(Fig. 1j-1l and Fig. S1f)**.

When genotyping complex SVs like inversions and duplications, the performance differences between the software are obvious. Although all software can detect inversion, VG-MAP, VG-Giraffe, GraphAligner, and Gramtools only worked effectively when the number of genomes was one. On the other hand, Paragraph and BayesTyper demonstrated superior performance with F-scores over or around 0.8 when genotyping inversion and duplications in graphs containing multiple *A. thaliana* or rice genomes **(Fig. S3a, S3b)**. Apparently, these tools presented lower F scores for genotyping heterozygous SVs, especially the duplications **(Fig. S3)**.

### Performance on plant genomes with different complexity

Moreover, tests conducted across various plant genomes revealed that certain tools, such as GraphAligner and GraphTyper2, exhibited relatively poor performance when dealing with larger genomes. Conversely, BayesTyper and PanGenie were able to retain high precision and recall even when working with more complex genomes **(Fig. 1 and Fig. S1)**.

As numerous plant genomes are heterozygous, we also simulated heterozygous *A. thaliana* and rice for the same test. Among the tools we evaluated, Paragraph, BayesTyper, and GraphTyper2 were less affected by heterozygosity, whereas other software experienced a decrease in recall for all variants **(Fig. S1 and Fig. S2).** We also explored the influence of heterozygosity on genotyping by testing on synthetic diploid genomes with varying levels of heterozygosity **(Fig. 2 and Fig. S4)**. The results showed that the damage to genotyping was proportional to the level of heterozygosity, especially for deletions and inversions. Paragraph and BayesTyper2 proved to be the most stable tools, both with high precision (0.75-0.97) and recall (0.79-0.96) for small and large indel genotyping **(Fig. 2 and Fig. S4)**. However, tools such as VG-MAP, VG-Giraffe, GraphAligner, and Gramtools were relatively more influenced by heterozygosity **(Fig. 2 and Fig. S4)**, especially for genotyping in repetitive regions **(Fig. S5b)**.

**Fig. 2.**
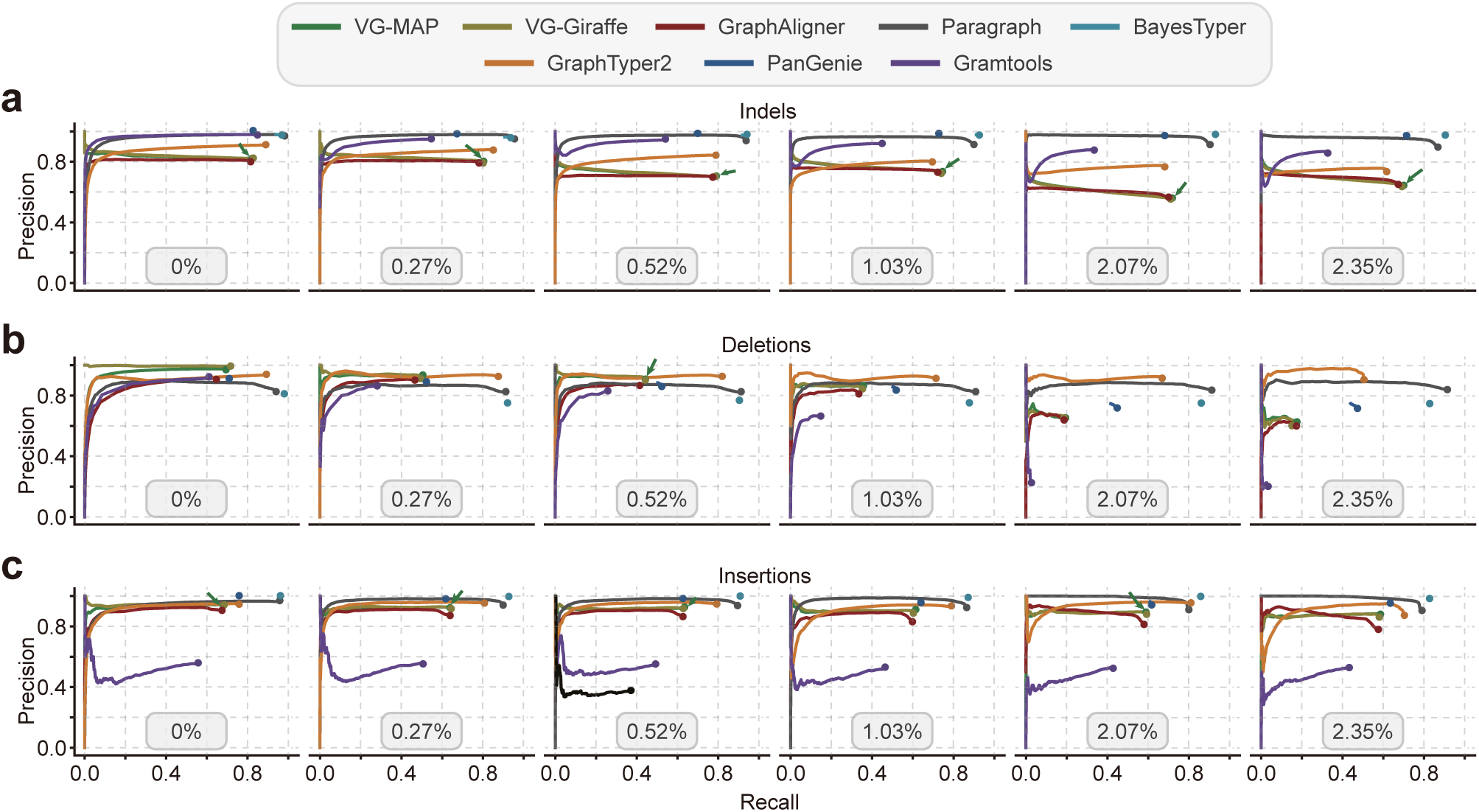
The effect of heterozygous rate on the performance of different graph-based genotyping methods. The six ROC curve plots correspond to the genotyping results for synthetic heterozygous *A. thaliana* genomes with different heterozygous rates (0%, 0.27%, 0.52%, 1.03%, 2.07%, and 2.35%). The genome graph for genotyping is constructed from the *A. thaliana* reference genome and seven alternative genomes. Paired-end (2×150 bp) short reads with 30× depth are simulated for genotyping. For each genotyper, precision is plotted against recall as the genotyping quality threshold varies. Read depth (DP) on variant sites is used as a proxy score when genotype quality (GQ) is not available. Arrows indicate the circles hidden by other circles in the plot due to identical or nearly identical precision values.

### Influence of sequencing parameters

Next, we conducted a comparison of each genotyper’s performance across datasets of paired-end reads with a range of read lengths (100 bp, 150 bp, 250 bp), fragment size (300 bp, 400 bp, 500 bp, 600 bp), and sequencing depths (5×, 10×, 20×, 30×, 50×). When the read length was shorter, such as 100 bp, only Paragraph showed a similarly high F-score for both small and large variants compared to testing with longer reads **(Fig. S6)**. Other tools, such as VG-Map and PanGenie, also had close F-scores with shorter (100 bp) or longer reads (150 bp, 250 bp) for variants except inversions. Additionally, a marginal effect could be observed when increasing read length from 150 bp to 250 bp **(Fig. S6)**. Apparently, various types of variants had similar requirements for sequencing length **(Fig. S6)**. The performance on rice and heterozygous *Arabidopsis* genomes followed the same trend as *A. thaliana*, suggesting that genome size and complexity may not affect the read length requirement by software **(Fig. S7-S8)**. In addition to using short reads, we also tested GraphAligner to map third-generation data against the genome graph, using long reads of 20kb and 75kb. The genotyping accuracy of 20kb reads was superior to that of 75,000 bp reads, likely due to the former’s higher accuracy (0.96 vs. 0.85) **(Fig. S6-S8)**. Furthermore, when the read length was 150 bp, small fragment sizes (300-600 bp) had no obvious effect on the genotyping accuracy of these genotypers **(Fig. S9)**.

Overall, when the sequencing depth was around 5-10×, Paragraph was able to achieve relatively high performance (precision > 0.81, recall > 0.91), whereas other software required more than 10× reads. **(Fig. S10)**. Besides, increasing the sequencing depth beyond 20×, only brought marginal improvements in genotyping performance across all variant types. With 30× data, all tested software almost reached the upper limit of genotyping precision and recall **(Fig. S10)**. These findings were consistent across different genomes, including the rice and heterozygous *A. thaliana* genomes, suggesting that genome size and complexity may not affect the sequencing depth requirements of these genotypers **(Fig. S11-S12)**. Besides, genotyping SVs requires more sequencing data than SNPs and indels **(Fig. S10-S12)**.

### Effects of genome number in the graph

As the search space of alignment may expand exponentially when more variants or genomes are graphed, we next evaluated how the number of graphed genomes affects variant genotyping. We reconstructed genome graphs for *A. thaliana* with a range (1, 7, 15, 30, 50) of individual genomes. From our evaluation, only tools Paragraph, BayesTyper, and GraphTyper2 demonstrated relatively stable precision and recall for SNP, indel, and insertion genotyping when graphed genomes increased **(Fig. 3a**, **3c and Fig. S13a)**. When only seven alternative genomes’ variants were graphed, existing methods could still achieve a good genotyping F-score (0.85-0.99 for SNPs, 0.81-0.97 for indels, 0.73-0.93 for deletions, 0.79-0.98 for insertions, 0.01-0.89 for inversions). However, when 50 alternative genomes were incorporated into the genome graph, the F-scores of deletion genotyping decreased to only 0.4-0.74. The recall of genotyping for VG-MAP, VG-Giraffe, and GraphAligner decreased as more genomes were incorporated. For example, as the number of graphed genomes increased from 1 to 50, the recall rate of VG-Giraffe decreased considerably (0.97 vs. 0.61 for SNPs, 0.99 vs. 0.66 for indels, 0.99 vs. 0.48 for deletions, and 0.92 vs. 0.53 for insertions, 0.55 vs. 0.23 for inversions), while the precision did not change much **(Fig. 3a and Fig. S13)**. In contrast, Paragraph, BayesTyper, and GraphTyper2 showed greater robustness in terms of recall **(Fig. 3a and Fig. S13)**. For example, BayesTyper’s recall rates remained above 0.95 for all types of variants except duplication (0.78), but the genotyping precision of deletion and inversion decreased by 0.52 (1.0 vs. 0.48) and 0.49 (1.0 vs. 0.51), respectively. Notably, the insertion precision of PanGenie was higher than that of deletion, possibly because PanGenie needs to count the number of k-mers for each haplotype at nodes. However, as only breakpoint sequences can be used for deletions, this will result in reduced genotyping precision.

**Fig. 3.**
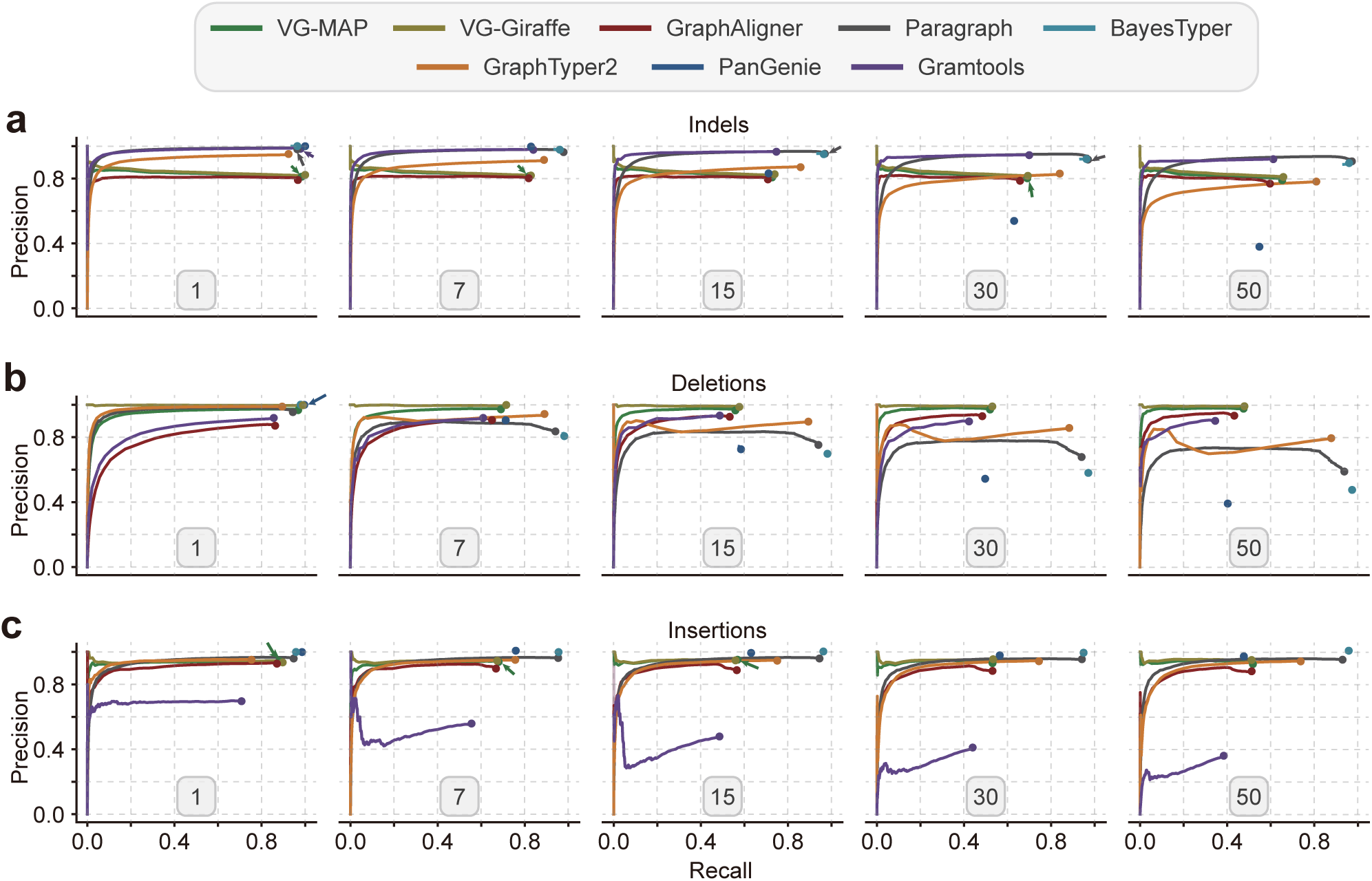
The effect of genome number on the performance of different graph-based genotyping methods. The five ROC curve plots correspond to genotyping results with different numbers (1, 7, 15, 30, 50) of graphed genomes. The genome graph for genotyping is constructed from the *A. thaliana* reference genome and different numbers of alternative genomes. Paired-end (2×150 bp) short reads with 30× depth are simulated for genotyping. For each genotyper, precision is plotted against recall as the genotyping quality threshold varies. Read depth (DP) on variant sites is used as a proxy score when genotype quality (GQ) is not available. Arrows indicate the circles hidden by other circles in the plot due to identical or nearly identical precision values.

### Influence of breakpoint deviations on SVs genotyping

Furthermore, we assessed the impact of breakpoint errors on SV genotyping by introducing a 1-20 bp deviation to the genuine breakpoints. For almost all tools, a negative correlation was observed between F-scores and breakpoint deviations **(Fig. 4a and Fig. S14).** Consistent with the previous report, BayesTyper, which is based on exact k-mer alignments, was more susceptible to breakpoint deviations. However, another k-mer alignment based genotyper, PanGenie, performed much better than BayesTyper. This may be because PanGenie can also leverage already known haplotypes to infer genotypes [19].

**Fig. 4.**
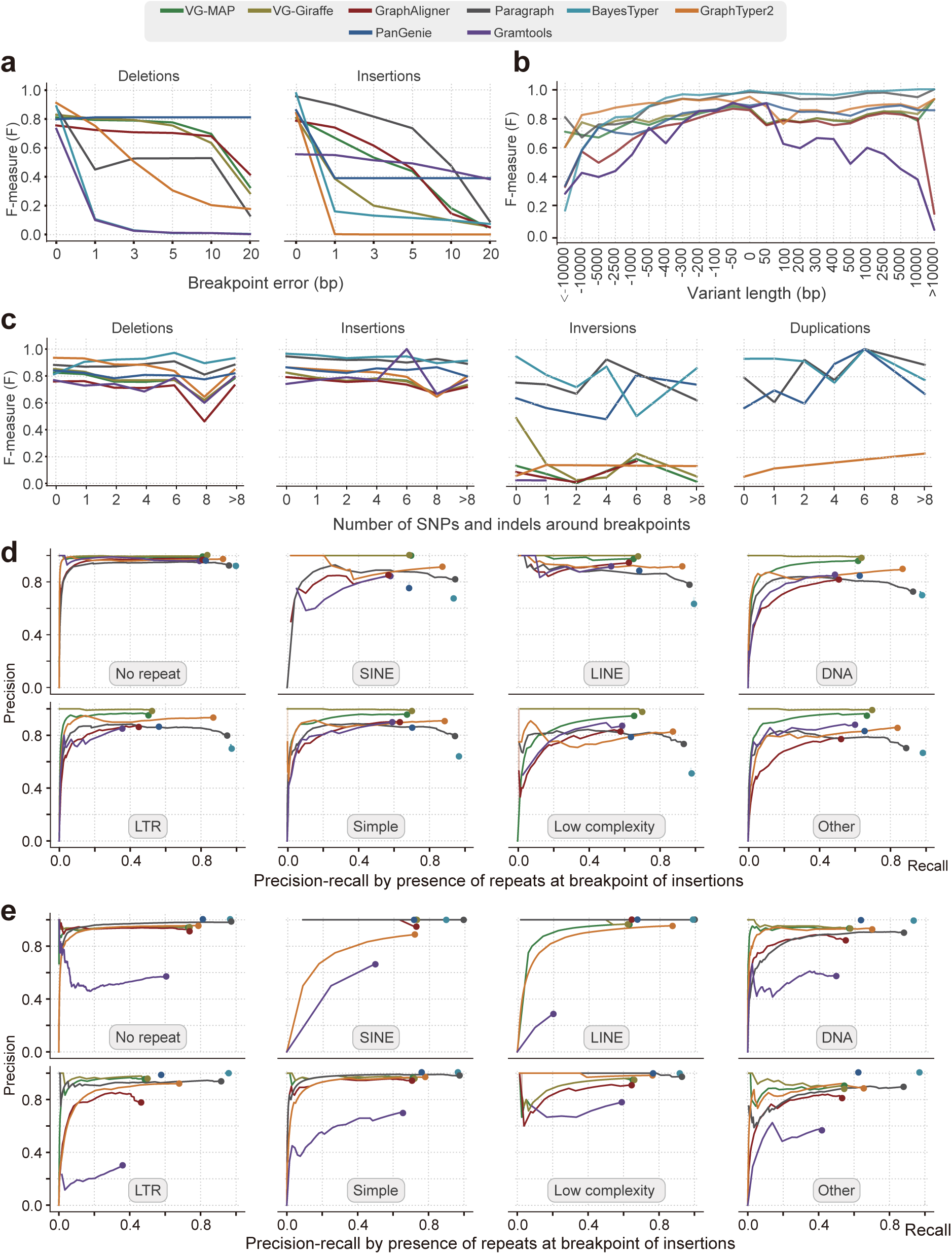
The impact of sequence context and event size on different graph-based genotyping methods. (a) Impact of breakpoint errors on genotyping (0 bp, 1 bp, 3 bp, 5 bp, 10 bp, 20 bp). **(b)** Impact of variant length (alternative allele length – reference allele length) on genotyping. **(c)** Impact of number of SNPs and indels around breakpoints on genotyping**. (d, e)** Effects of repeat sequence types around breakpoints on genotype, partitioned by variant type: (d) deletions and (e) insertions. Different types of repeat sequences are annotated by RepeatMasker. The genome graph for genotyping is constructed from the *A. thaliana* reference genome and seven alternative genomes. Paired-end (2×150 bp) short reads with 30× depth are simulated for genotyping. For each genotyper, precision is plotted against recall as the genotyping quality threshold varies. Read depth (DP) on variant sites is used as a proxy score when genotype quality (GQ) is not available.

Overall, the performance on genotyping deletion was better than that of insertion, inversion, and duplication. For genotyping deletion, PanGenie, VG-MAP, VG-Giraffe, Paragraph, and GraphAligner displayed consistent performance with breakpoint deviations smaller than 10 bp **(Fig. 4a)**. For genotyping insertions, the F-scores of VG-MAP, GraphAligner, Paragraph, and Gramtools still had 50% to 90% of that of 0 bp when the breakpoint error reached 5 bp, while the F-scores of other software at 1 bp deviation were reduced by more than 50% **(Fig. 4a)**. Further, only Paragraph and VG-Giraffe maintained performance within a 10 bp error on the genotyping of inversion **(Fig. S14a)**. However, none of the software was able to tolerate break-point errors in duplications **(Fig. S14b)**.

### Impact of event size and sequence context on SV genotyping

In addition, we stratified SVs based on event size, number of SNPs and indels within break-points of 100 bp, and families of repetitive sequences around to estimate the effect of sequence context on SV genotyping. Overall, Paragraph and BayesTyper demonstrated the best genotyping performance across different size ranges of SVs. Although the F-scores of both Paragraph and BayesTyper were lower (ranging from 0.17 to 0.81) for deletions larger than 5 kb, they still maintained high F-scores (≥ 0.95) for insertion **(Fig. 4b)**. Previous studies have reported that small variants near the breakpoints could affect the accuracy of SV calling [11]. However, our experiment showed that small variants had no serious damage on SV genotyping for these graph-based tools **(Fig. 4c)**, which might be attributed to the improved alignment accuracy as alternative alleles (small variants) are introduced into the graphs.

The genotyping performance of all methods was reduced in the repetitive regions compared to the nonrepetitive regions **(Fig. 4d**, **4e and Fig. S13)**. For example, the GraphAligner F-score was even 47% lower in deletions than non-repetitive regions **(Fig. S5)**. Compared to other software, Paragraph and BayesTyper demonstrated comparatively stable performance in repetitive regions **(Fig. S5)**. Repeat sequences had a more severe influence on the genotyping of deletions (17%) than SNPs (9%), indels (11%) and insertions (8%) **(Fig. 4d**, **4e and Fig. S13a)**. Similar to the linear reference-based SV genotyping [11], LTRs had the greatest impact on genotyping. For example, the recalls of deletion genotyping for VG-MAP, VG-Giraffe, and GraphAligner were reduced by 0.31, 0.31, and 0.36, respectively **(Figs. 4d and 4e)**.

### Development of an ensemble genotyper

In summary, these graph-based genotypers performed differentially for small and large variants in terms of precision and recall. Further analysis on the overlap of true variant genotyping among the eight genotypers revealed that many variants were not correctly genotyped by some genotypers but were correctly genotyped by others **(Fig. S15,** example shown in **Fig. S16)**, suggesting an ensemble genotyping strategy may improve genotyping performance.

Here, we developed an Ensemble Variant Genotyper (EVG) by integrating various graph-based genotyping methods **(Fig. 5a)**. Before running the genotyping pipeline, EVG modifies VCF-formatted variants input files provided by users to a common format that can be used for downstream analysis (EVG convert). Additionally, EVG could filter variants based on minor allele frequency (MAF) and missing rate, if indicated. The EVG pipeline starts with the selection of the most suitable software according to the reference genome size, sequencing data quality, type and number of variants, and software preference. The selected software is then run, respectively. To address the issue caused by inconsistent coordinates of the same variants from different software, EVG then clusters the outputs based on the size and position of variants and constructs a variant graph (EVG merge). Finally, the most probable combination of genotypes is determined by calculating the number of reads supporting each genotype at each node **(Fig. 5a)**. To accelerate genotyping, EVG can randomly downsample reads when sequencing data is high enough. For oversized genome graphs with numerous variants, EVG offers an optional fast mode where specific software is used only for particular types of variants. These allow EVG to significantly accelerate genotyping while sacrificing very little precision and recall **(Fig. 5b)**.

**Fig. 5.**
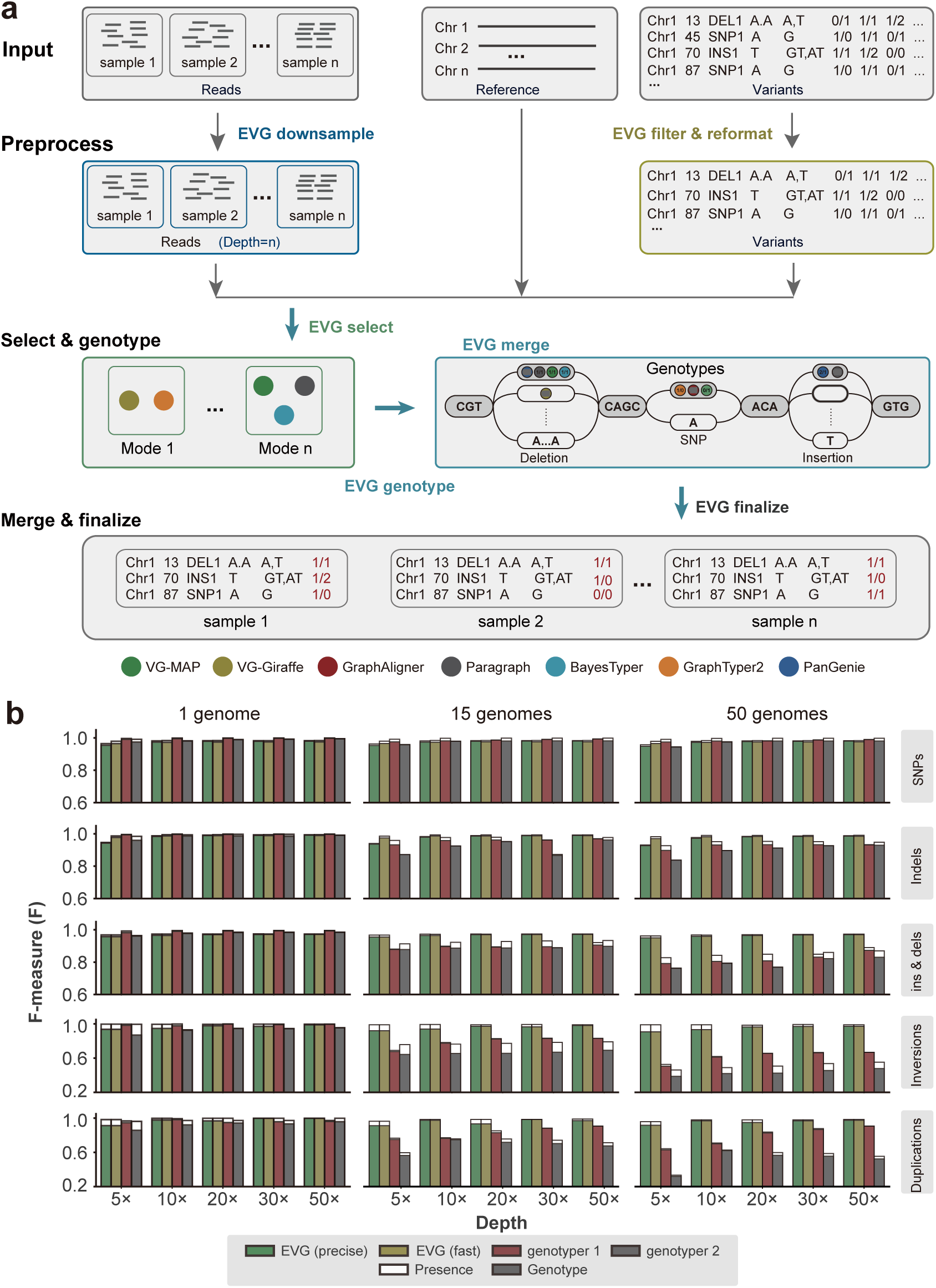
The workflow and performance of the ensemble variant genotyping method EVG. (a) The variant genotyping workflow of EVG mainly consists of three steps: 1) subsample sequencing reads, filter variants, and reformat the input variant VCF file; 2) select one or multiple suitable graph-based genotypers (shown as colored dots) and do genotyping with each of them in parallel; 3) merge the genotype results from step 2 and determine the final genotype for each variant. **(b)** Genotyping performance of SNPs, indels, ins & del (insertions and deletions), inversions, and duplications on simulated *A. thaliana* genomes under different sequencing depths (5×, 10×, 20×, 30×, 50×) and genome numbers (1, 15, 50). The genome graph for genotyping is constructed from the *A. thaliana* reference genome and different numbers of simulated alternative genomes. Paired-end short-reads (read length: 2×150 bp) are simulated for variant genotyping. For each genotyping scenario, the F-measure values of the other two best-performing genotypers are shown here.

To assess the performance of EVG, we tested it on all simulated datasets from this study. Firstly, unlike other graph-based genotypers, EVG achieved the highest F-score for both small and large variants with just 5× 150bp paired-end short-reads **(Fig. 5b and Fig. S17)**. Secondly, EVG’s performance was more robust when more genomes were graphed. Specifically, for genome graphs with 50 genomes, EVG achieved a F-score above 0.95 for SVs with only 5× short reads, while other best genotypers only reached 0.79 **(Fig. 5b and Fig. S17)**. In terms of SNP genotyping, the fast mode of EVG performed slightly lower than the best-performing software because, to reduce CPU time usage, EVG only chose two software for SNP genotyping **(Fig. 5b)**.

### Performance on real data

Finally, we carried out testing on real data consisting of 30× Illumina short reads from three diploid homozygous genomes (*A. thaliana*, rice and maize) [5, 38, 40, 42, 43] as well as one diploid heterozygous genome of Apricot (*Prunus armeniaca*) [44, 45]. Testing was also performed on a genome graph of seven genomes for all genotypers. **(Table S5)**. Notably, Gramtools was also excluded from the real data testing. In comparison to the simulated data, the F-score across genotypers in the real data was lower (**Fig. 6**). Such a reduction was even worse (< 0.65) for the maize genome, probably because of the high percent of repetitive sequences **(Fig. S18)**. For the heterozygous apricot genome, Paragraph had the highest average F-score in all types of variants compared to other six graph-based genotypers, while BayesTyper was hardly able to genotype the SVs, perhaps due to inaccurate breakpoints **(Fig. 6)**.

**Fig. 6.**
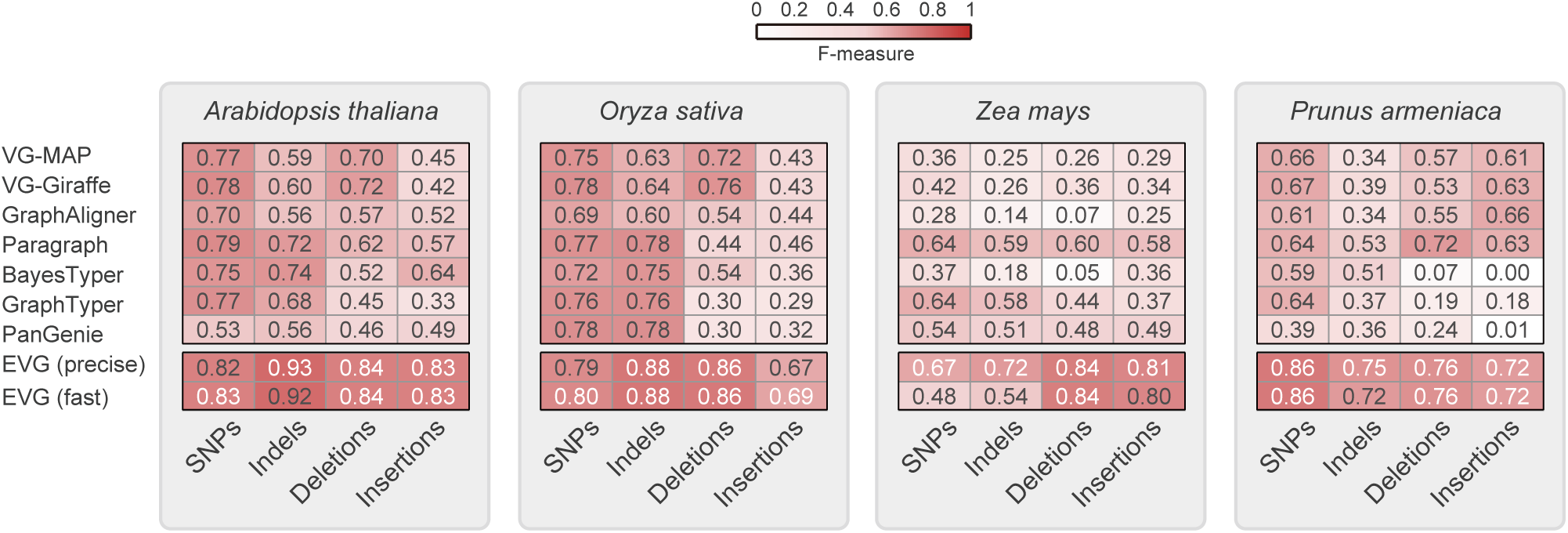
Overall genotyping performance of different graph-based methods based on real data. The read lengths of rice (*Oryza Sativa*), maize (*Zea mays*), and apricot (*Prunus armeniaca*) are all 2×150 bp, except for *A. thaliana* (2×100 bp). The sequencing depth and number of graphed alternative genomes are 30× and 7, respectively.

We also tested EVG’s performance on all real data sets and found that for all four genomes, EVG achieved the best genotyping performance across all types of variants. Notably, for the largest genome, maize, EVG (precise model) reached the highest F-score (0.67 for SNPs, 0.72 for indels, 0.84 for deletions, and 0.81 for insertions) **(Fig. 6)**. Similarly, for the heterozygous genome, EVG showed a comparably higher F-score ranging from 0.72 to 0.86 for genotyping small and large variants. More importantly, EVG’s performance in repeated regions was also better than other software, especially for deletion (F-score, 0.82) and insertion (F-score, 0.83) **(Fig. S18)**. In summary, our pipeline can be applied to plant genomes with a range of genome sizes or repeat contents compared to the currently available graph-based variant genotypers.

### Runtime and memory usage

The runtime and memory usage of all the methods were measured with the same number of CPUs. As expected, read alignment-based methods such as VG-MAP and Paragraph require relatively more time compared to k-mer alignment-based methods, which were faster despite having larger memory consumption **(Fig. S19a-S19c)**. Additionally, runtime and/or memory increase significantly when graphing more or larger genomes. For instance, VG-MAP’s runtime in *A. thaliana* was over 150 times that in maize **(Fig. S19)**. GraphTyper2 required low runtime and memory usage for smaller genomes like *A. thaliana* and rice, and slightly more runtime for maize. When tested under the same conditions, EVG’s fast mode required only 5.5 CPU hours in *A. thaliana* and 21.6 CPU hours in rice, and its genotyping was more robust than that of existing methods. The precise mode of EVG further improved the genotyping performance, but it took 16.7 and 63.8 CPU hours in *A. thaliana* and rice. In general, EVG has a comparable running time to VG-Giraffe and k-mers-based software, but its genotyping performance is superior to other software **(Fig. S19-S20)**.

## Discussion

In this study, we have conducted a comprehensive evaluation of the performance of eight popular graph-based genotypers on multiple representative plant genomes, which are mostly designed and tested only on human genomes. By conducting tests on 25 simulation and 10 real datasets, we have revealed the differences in precision and recall among these tools under different sequencing schemes, genomic context and complexity, and graph size for a spectrum of genetic variants. More importantly, the EVG pipeline developed here can achieve comparably higher genotyping recall and precision even when using 5× short reads and remain stable with increased number of genomes, fitting the trend requiring population-level millions of variant genotyping.

Plant genomes frequently have a large genome size, enriched repetitive sequences, and high sequence diversity or heterozygosity [36, 37]. These features seriously challenge the efficiency of constructing and indexing pangenome graphs and read alignments. For example, graphing seven maize genomes with PanGenie, BayesTyper, and Paragraph requires over 100 GB memory **(Fig. S15)**. To address these challenges, BayesTyper genotypes local haplotypes by constructing a Bloom filter for selecting which read k-mers should be loaded into memory (i.e., those k-mers stored in the Bloom filter) [18]. However, storing k-mer counts and variant graphs in memory may lead to significant memory consumption, particularly when dealing with large amounts of sequencing data or reference genomes. Alternatively, implementing a Counting Bloom filter or Hierarchical Interleaved Bloom Filter (HIBF) instead of storing k-mer counts in memory may help reduce memory consumption while maintaining the desired level of accuracy [46, 47]. PanGenie, another software program employing k-mer counts, exhibits significant memory consumption despite only considering unique k-mers [19]. While its memory footprint exceeds that of BayesTyper in *A. thaliana*, it is comparatively lower in rice and maize. This phenomenon may be attributed to the abundance of repeated k-mers in these plant species that are filtered out during the analysis process.

Unlike k-mer alignment based genotypers, the VG tool utilizes less memory during genome graph construction and relies on global read alignments [16]. However, the earlier version, VG-MAP requires a significant amount of memory and CPU time for graph indexing, particularly when constructing GCSA2 indexes [34]. To address these issues, VG-Giraffe [20] (and GraphAligner [28]), use minimizers [35] for seeding to speed up the alignment and reduce memory consumption at the expense of the alignment rate for repeat sequences. In contrast to VG and GraphAligner, GraphTyper2 [17] and Paragraph [24] utilize a local variant realignment strategy based on pre-aligned reads, or solely mapping those around breakpoints. Although such an approach theoretically has the potential to markedly accelerate the genotyping process, our testing revealed that the runtime of Paragraph did not demonstrate significant improvement. This can be attributed primarily to the I/O overhead as three files are generated for each variant during the genotyping. Therefore, despite recent advances in genome graph software, the high variability of plant genomes remains a major challenge for current approaches.

Additionally, SV breakpoints often exhibit deviations [10], particularly at repeat-enriched regions, which can affect some genotypers [24], especially those such as BayesTyper, that require long exact matches of k-mers or seeds for read mapping. However, another k-mer based genotyper, PanGenie, performs much better than BayesTyper by integrating haplotype-resolved pan-genome references. The enriched heterozygous alleles and repetitive sequences will also affect the genotype effect of tools.

Most current graph-based genotypers require 10-20× of data to achieve satisfactory genotyping performance. However, the sequencing depth of early or large genome population sequencing projects in plants is low [39, 48], even only 3-10× for wheat [49]. Future development should consider how to distinguish between sequence errors and real variants especially those in heterozygous loci and repetitive regions with low sequence depth of short reads. One potential approach similar to that implemented in PanGenie is to integrate the known haplotypes of more haplotype-resolved genome assemblies. Additionally, these tools lack stability for population-level pangenome graph-based genotyping. This study found that as the number of variants increased, the performance of the software continuously decreased while the runtime and memory increased considerably.

Taken together, existing methods either require excessive computational resources or sequencing costs, or lack stable performance for plant genomes of different complexity, which limit their applicability in plants. To alleviate these problems, we have developed the EVG pipeline, which select the most appropriate process based on reference genome characteristics, sequencing data, heterozygous rate, and read depth. And the fast mode of EVG will provide specific variants to specific genotypers according to their preferences. EVG pipeline presents more robust genotyping performance compared to existing graph genome methods. More importantly, EVG can reach high F-scores of genotyping with only 5× reads, and achieve the peaks with 10× reads, which is normally the average depth of population-level resequencing in plants. Moreover, even with an increased number of nodes in the graph (up to 50 genomes), the genotyping F-score of EVG remains above 0.9. **(Fig. 5b)**. Most notably on maize, the final genotyping results were better than those obtained using other software, requiring only 210 CPU hours (including the time of graph construction, graph indexing, read alignment, and genotyping, 2666 CPU hours for VG-MAP) **(Fig. S19)**. EVG also performs exceptionally well in maize repetitive regions, achieving F-scores for deletions and insertions of 0.82 and 0.83 respectively, while the highest values of other software were 0.59 and 0.55.

Although the EVG pipeline has several advantages compared to current graph-based genotypers, it can still be improved in terms of memory consumption and genotyping small variants in the future. The further application of graph-based genotyping in plants requires more consideration for complex regions with dense variants, highly similar regions due to whole genome duplication, and polyploidy genomes. As more high-quality genomes and variants are obtained by long read sequencing technologies, population-level and type-full variant genotyping with short reads will be practicable by using graph-based methods, thus facilitating population analysis or trait association studies. By comprehensively testing multiple plant genomes, we reveal the performance level of these graph-based genotypers in different scenarios. Our EVG pipeline with higher performance and stability should be applied to population-scale genotyping for millions of all types of genetic variations for genomes with lower sequencing costs.

## Conclusions

This paper comprehensively evaluates multiple genome graph-based genotyping software packages using both simulated and real data sets. The results reveal significant challenges in applying existing genome graph software to plants, including resource-intensive computing, poor genotyping accuracy for repeat-related variants, and unstable genotyping performance. The EVG pipeline developed here delivers superior genotyping performance even in repeat regions with minimal increases in resource consumption when only 5× short reads are provided. Our EVG pipeline will be potentially used in population-scale variant genotyping and contribute to plant pan-genomic research.

## Methods

### Selection of graph-based genotyper

The following genotypers were selected: VG v1.37.0, GraphAligner v1.0.13, Paragraph v2.3, BayesTyper v1.5, GraphTyper2 v2.7.2, PanGenie v2.0.0, and Gramtools v1.10.0. Both VG-MAP and VG-Giraffe are used for read alignment in the VG package.

### Simulated datasets

Overall, the simulated data include sequencing reads from different genomes (*A. thaliana*, rice, maize, *Brassica napus,* heterozygous *A. thaliana*, and heterozygous rice) with different sequencing parameters (technologies, read length, fragment size, and sequencing depth) **(Table S3 and S4)**. The sequencing reads were used for genotyping variants from public variation databases and/or resulting from genome comparisons.

We used the ART [50] software (version 2.1.8) to generate Illumina paired-end short reads for each of the alternative genomes derived by introducing variants into the reference genomes using the VarSim [51] (version 0.8.6) simulator. For *A. thaliana*, variants from the 1001 Genomes Project and one from our previous study [5, 38] and the reference Col-0 from TAIR10 (https://www.arabidopsis.org) [52] were used. For rice, all types of variations from the Rice SNP-Seed Database (https://snp-seek.irri.org/) [53] and the reference IRGSAP-1.0 (https://rapdb.dna.affrc.go.jp/) [54] were used. For maize, variants are from whole genome comparisons between the reference B73 v5.0 (https://www.maizegdb.org) [40, 43] and previously released assemblies of different accessions [40, 43]. These assemblies were aligned to the reference using Minimap2 [55]. Show-snps from MUMmer4 [56] was used for calling SNPs and indels, and Assemblytics [57] was used for calling SVs. For *Brassica napus*, variants from whole genome comparisons between the reference assembly ZS11 (http://cbi.hzau.edu.cn/bnapus/) [41] and previously released assemblies of different accessions [41]. The method used for genome comparisons and variant callings were the same as in maize.

The number of introduced variants **(Table S4)** is similar to the average number of variants found in real *A. thaliana* accessions [38]. The same control of variant numbers was done for other plant genomes. To evaluate the genotyping when multiple genomes are graphed, we also introduced more variants obtained from the databases described above to simulate multiple genomes **(Table S4)**.

To evaluate the genotypers’ performance on heterozygous genomes, we simulated heterozygous genomes for *A. thaliana* and rice. Because VarSim cannot specify the degree of heterozygosity, genome heterozygosity can only be controlled by adjusting the number of variants and the percentage of heterozygous variants (with parameters of ‘vc_prop_het’ and ‘-- sv_prop_het’,). Finally, five genomes with different heterozygosity rates (0.27%, 0.52%, 1.03%, 2.07%, and 2.35%) were simulated.

To evaluate the genotypers’ performance under different sequencing parameters, we used the ART simulator to simulate short paired-end reads with a range of read length (2×100 bp, 2×150bp, and 2×250 bp), fragment size (300 bp, 400 bp, 500 bp, 600 bp) and sequencing depth (5×, 10×, 20×, 30× and 50×). Simulated PacBio sequencing (P6C4 model) was generated using PBSIM [58] (version 2.0.1) with the simulated *A. thaliana* genome serving as the reference. Varying read lengths (20kb and 75kb bp) and accuracies (0.96 for 20kb and 0.85 for 75kb bp) were generated.

### Real datasets

*A. thaliana* real datasets were from the 1001 Genomes Project [5] and one previous study (with NCBI project ID PRJEB31147 [38]), including genome sequences, PacBio, and Illumina sequencing reads (accessions: An-1, C24, and Cvi). Rice real datasets were from the study Zhang et al. [42], downloaded from the CNCB (project ID: CRA004623) and TGSrice databases, including genomes, ONT, and Illumina sequencing reads (accessions: TG19, TG28 and TG78). Maize real datasets were from the study Hufford et al [43], downloaded from the CNCB (project ID: PRJEB31061) and MaizeGDB databases [40], including genomes, ONT, and Illumina sequencing reads accessions: B97, CML52, and CML69).

To construct the benchmark variant dataset, we first map short reads with bwa [59] (version 0.7.17-r1198-dirty) and call SNPs and indels with GATK [60] (version 4.2.6.9) and BCFtools [61] (version 1.9). We kept variants shared by the two tools, quality score larger than 20 and a sequencing depth lower than 100. For the SV dataset, we mapped long reads using NGMLR [62] (version 0.2.7) with default parameters and subsequently detected SVs using Sniffles [62] (version 2.0.3). In addition, we utilized the nucmer tool [56] (version 4.0.0rc1, parameters: ‘-c 100’, ‘-b 500’, and ‘-l 50’) to align the alternative genome against the reference genome and subsequently identified SVs using Assemblytics [57] (version 1.2.1, parameters: unique sequence length of 10,000, minimum variant size of 50, and maximum variant size of 100,000). We identified those SV common between Sniffles and Assemblytics by filtering those with breakpoint differences more than 200 bp or event size differences larger than 25% of the real event size. The resulting SVs shared by Sniffles and Assemblytics were used for genotyping.

For genotyping evaluation on heterozygous genomes, we used one haplotype-resolved and chromosome-level assembly of apricot (*P. armeniaca*; cultivar “Rojo Pasión”) from one previous study [44]. The Illumina short reads and PacBio long reads from this study were also used for building the variant dataset. The reference genome from cultivar “Yinxiangbai” is used for the read mapping [45]. Variants shared by the two haplotypes are homozygous, and the specific ones are heterozygous. To obtain a high-quality variant dataset, we applied both read mapping and assembly comparison based methods. Firstly, SNPs and indels were called by GATK and BCFtools based on short read mapping, and those common ones were retained, similar to what was described above. Secondly, SVs were called by Sniffles based on the PacBio read alignments resulting from NGMLR. Thirdly, the two haplotype assemblies were aligned to the reference by nucmer (parameters: ‘-c 100’, ‘-b 500’, and ‘-l 50’), followed by calling SNPs and indels with show-snps, and SV with Assemblytics (parameters: unique sequence length of 10,000, minimum variant size of 50, and maximum variant size of 100,000). Finally, variants shared by the read mapping method and the assembly comparison method were identified using the same criteria with other genomes as described above.

### Variants genotyping with simulated and real datasets

For genotyping with simulated homozygous *A. thaliana* and rice genomes, different numbers of genomes (1, 7, 15, 30, and 50) were graphed. For genotyping with simulated heterozygous *A. thaliana* and rice, graphs with one and seven genomes were used. For genotyping with simulated maize and *Brassica napus*, graphs with one and seven genomes were used. For genotyping with real datasets from *A. thaliana*, rice, maize, and apricot, graphs with one and seven genomes were used. For genotyping evaluation in a real dataset from *A. thaliana*, rice, and maize, variants from three different accessions were genotyped individually. The detailed information on genome graphs and variants is included in **Table S4-S6**.

As genotypers Paragraph, BayesTyper, and GraphTyper2 require linear-reference based read alignment BAM files, we used BWA to align paired-end short reads from each dataset and used SAMTools [8] (version 1.15) to sort and convert the alignment output into BAM format. VG-MAP, VG-Giraffe, GraphAligner, PanGenie, and Gramtools all directly input read data, variant data, and reference genomes. For all software except GraphTyper2, we run the genotyping with the default parameters as recommended in their manuals. The detailed commands for running each genotyping tool on a Linux system are uploaded to the GitHub website (https://github.com/JiaoLab2021/EVG/wiki/EVG-paper).

To measure the genotyping precision, recall, and F-score, we compared each variant called by each genotyper with the true variant dataset by using the script graphvcf in EVG developed in this study. For SVs, if both the start breakpoint and the end breakpoint of one SV were within 200 bp of the true SV positions, and the SV sizes differed by at most 25% of the true size, such SV calling was considered correct presence. If the genotype of this SV calling also matched the true event, it is considered correct genotype. For an indel calling, the correct presence requires a position difference less than 10 bp, while for a SNP calling, an exact position match is necessary.

### The EVG workflow

EVG software takes as input a variant VCF file of the population, the reference genome, and a configuration file containing the sequencing reads path. The whole EVG workflow mainly contains three steps. EVG can support restarting the task from the point of failure by using the ‘--restart’ parameter. The details of each step are described as follows:

#### Step 1: preprocessing

This preprocessing step mainly involves the read data subsampling, variant filtering, and VCF file reformatting. Firstly, to speed up read mapping against genome graphs, EVG offers a solution by first extracting a subset (default: 15×) of read data for downstream genotyping. Based on this study, acceptable genotyping results can be achieved with as little as 5× data. Secondly, to reduce resource consumption, variants can be filtered according to their Minor Allele Frequency (MAF) and missing rate using EVG. By default, variants are not filtered. Thirdly, to avoid throwing errors due to incompatible input variants in the VCF file provided by users, EVG automatically checks the VCF file and reformats any variants to be compatible with the software’s different requirements accordingly.

#### Step 2: select genotypers and do genotyping

EVG automatically selects the optimal genotyping process based on factors including the size of the reference genome, the sequencing depth of the individual genome to be genotyped, and the read length of the sequencing data. According to our evaluation results, Paragraph and BayesTyper relatively perform better in terms of accuracy, recall, runtime, and memory consumption. Thus, these two genotypers are both included in the EVG workflow. EVG adds the third genotyper, VG-MAP, while it adds VG-Giraffe instead when the genome is larger than 1Gb. Additionally, when the read length is greater than 130bp and the sequencing depth is greater than 5×, the genotyper GraphTyper2 is also included. If none of the previous steps include GraphTyper2, then PanGenie is used **(Fig. S21)**. Remarkably, Paragraph generates three files for each variant during running, which will result in high disk I/O consumption if SNPs and indels are included. EVG provides a ‘fast’ model where only variants larger than 50 bp will be genotyped by Paragraph. When the sequencing data is third-generation, EVG only uses GraphAligner. After selecting, EVG submits all tasks in parallel. When all tasks are completed, EVG converts the output for subsequent merges.

#### Step 3: merge and finalize genotyping

EVG will first cluster all the variants to form a variant graph, with each node containing the variant’s location, length, variant type, genotyping, and depth information. The variants in the same position are put on the same branch. To keep the read depth of different software on the same order of magnitude, we use the Z-score to normalize the read depth for each variant:

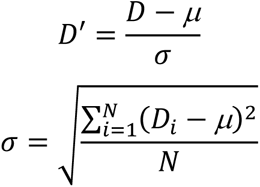

where *D’* indicates the normalized read depth of a variant. *D* corresponds to this variant read depth calculated by genotypers and present in the genotyping output VCF file. *μ* and *σ* are the average and standard deviation of read depth for all variants genotyped by a genotyper. *N* is the number of variants.

EVG selects the most likely genotyping according to consistency across different genotypers and depth information. For small variants, including SNPs and indels, the genotype is the most commonly used by genotypers. When no genotype is supported by more than one genotyper, EVG will skip this variant. For SV genotyping, the final genotype is from BayesTyper or Paragraph when their genotyping results are available (if both available, BayesTyper’s genotype is preferred), as these two genotypers presented relatively better performance in our evaluation. If neither of them has genotyping result for a given SV, the genotype is one from another genotyper with the smallest normalized absolute depth.

## Declarations

### Ethics approval and consent to participate

Not applicable.

### Consent for publication

Not applicable.

### Availability of data and materials

The source code of EVG, as well as the commands used to conduct the analyses presented in this study, are available on GitHub (https://github.com/JiaoLab2021/EVG/) under an MIT license. The whole genome sequencing data for *A. thaliana*, rice, maize, and apricot was downloaded from SRA (PRJEB31147 [38]), CNCB (CRA004623 [42], PRJEB31061 [43]), BioProject (PRJNA577047 [44]), ENA (PRJEB37669 [44]), and CNGBdb (CNP0000718 [44]).

### Authors’ contributions

WBJ conceived and designed the project. ZZD and JBH collected the data. ZZD and JBH conducted the analyses. ZZD developed the EVG package. ZZD and WBJ wrote the paper. All authors read and approved the final manuscript.

## Acknowledgements

We thank the high-performance computing platform at the National Key Laboratory of Crop Genetic Improvement at Huazhong Agricultural University.

