## Supplementary material for "A comprehensive benchmark of graph-based genetic variant genotyping algorithms on plant genomes for creating an accurate ensemble pipeline": Table S, Figure S

### Supplementary figures and tables

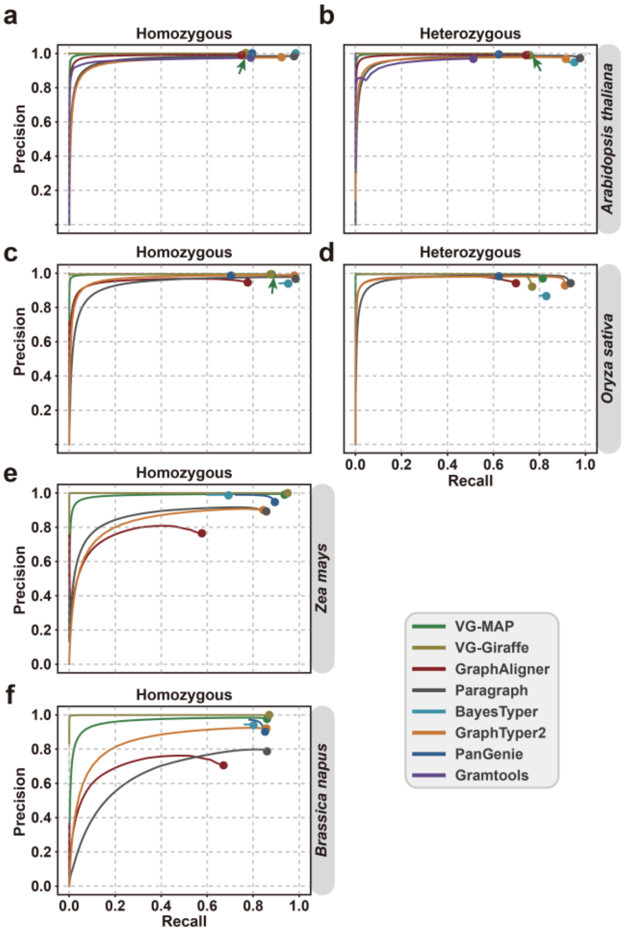


**Figure S1. Overall performance of SNP genotyping for different graph-based tools based on simulated data.** The genome graphs of *Arabidopsis thaliana* **(a, b)**, *Oryza sativa* **(c, d)**, *Zea mays*, **(e)** and *Brassica napus* **(f)** are constructed based on one reference genome and seven alternative genomes derived by introducing known variants into the reference genome. For *A. thaliana* and *O. sativa*, SNP genotyping performance on simulated heterozygous genomes were also evaluated. Paired-end (2×150 bp) short reads with 30× depth were simulated for genotyping. For each genotyper, precision is plotted against recall as the genotyping quality threshold varies. Read depth (DP) on variant sites is used as a proxy score when genotype quality (GQ) is not available. Arrows indicate the circles hidden by other circles in the plot due to identical or nearly identical precision values.

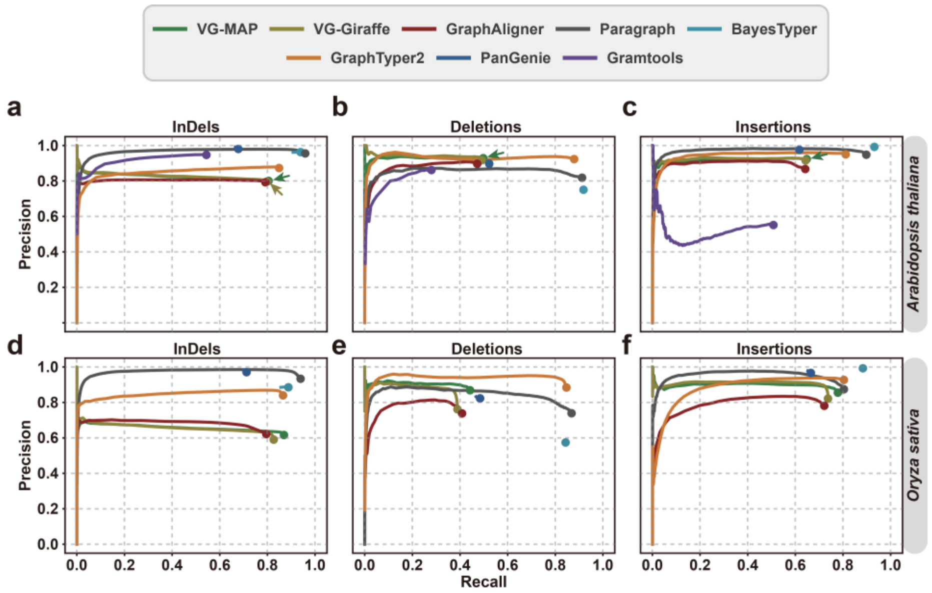


**Figure S2. Overall genotyping performance of different methods on heterozygous *Arabidopsis* and rice genome.** Variations are divided by type: **(a, d)** indels, **(b, e)** deletions, **(c, f)** insertions. The genome graphs of *Arabidopsis thaliana* **(a, b, c)** and *Oryza sativa* **(d, e, f)** are constructed based on one reference genome and seven alternative genomes derived by introducing known variants into the reference genome. Paired-end (2×150 bp) short reads with 30× depth were simulated for genotyping. For each genotyper, precision is plotted against recall as the genotyping quality threshold varies. Read depth (DP) on variant sites is used as a proxy score when genotype quality (GQ) is not available. Arrows indicate the circles hidden by other circles in the plot due to identical or nearly identical precision values.

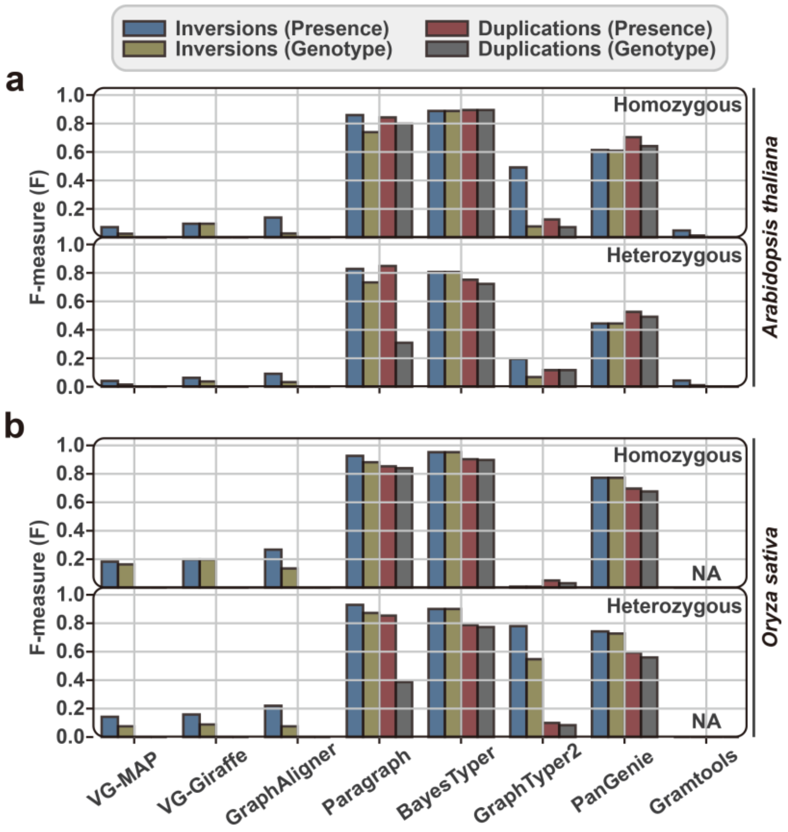


**Figure S3. Genotyping performance of inversions and duplications on simulated data. (a)** Genotyping performance of inversion and duplication in homozygous and heterozygous *Arabidopsis* thaliana. **(b)** Genotyping performance of inversions and duplications in homozygous and heterozygous Rice. The genome graphs of *Arabidopsis thaliana* **(a)** and *Oryza sativa* **(b)** are constructed based on one reference genome and seven alternative genomes derived by introducing known variants into the reference genome. Paired-end (2×150bp) short reads with 30× depth were simulated for genotyping.

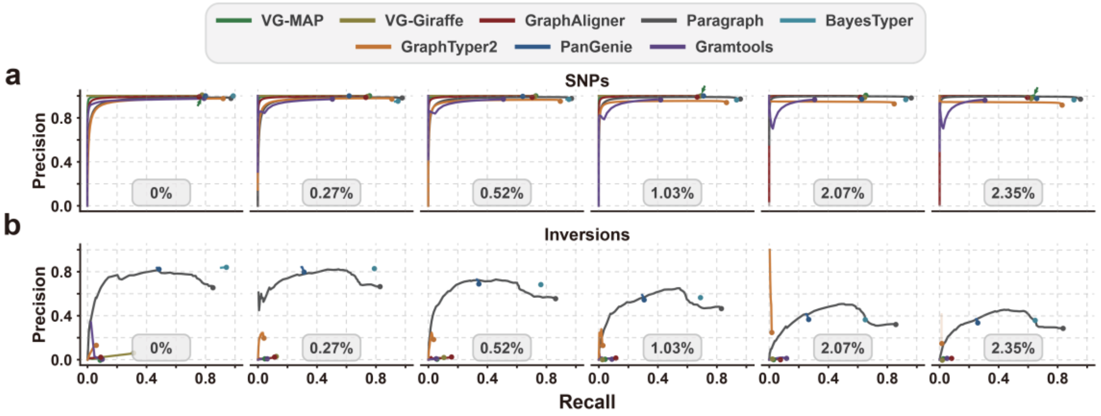


**Figure S4. The effect of heterozygous rate on the genotyping performance of different methods, partitioned by variant type: (a) SNPs, (b) inversions.** The six ROC curve plots correspond to the genotyping results for synthetic heterozygous *A. thaliana* genomes with different heterozygous rates (0%, 0.27%, 0.52%, 1.03%, 2.07%, 2.35%). The genome graph for genotyping is constructed from the *A. thaliana* reference genome and seven alternative genomes. Paired-end (2×150 bp) short reads with 30× depth are simulated for genotyping. For each genotyper, precision is plotted against recall as the genotyping quality threshold varies. Read depth (DP) on variant sites is used as a proxy score when genotype quality (GQ) is not available. Arrows indicate the circles hidden by other circles in the plot due to identical or nearly identical precision values.

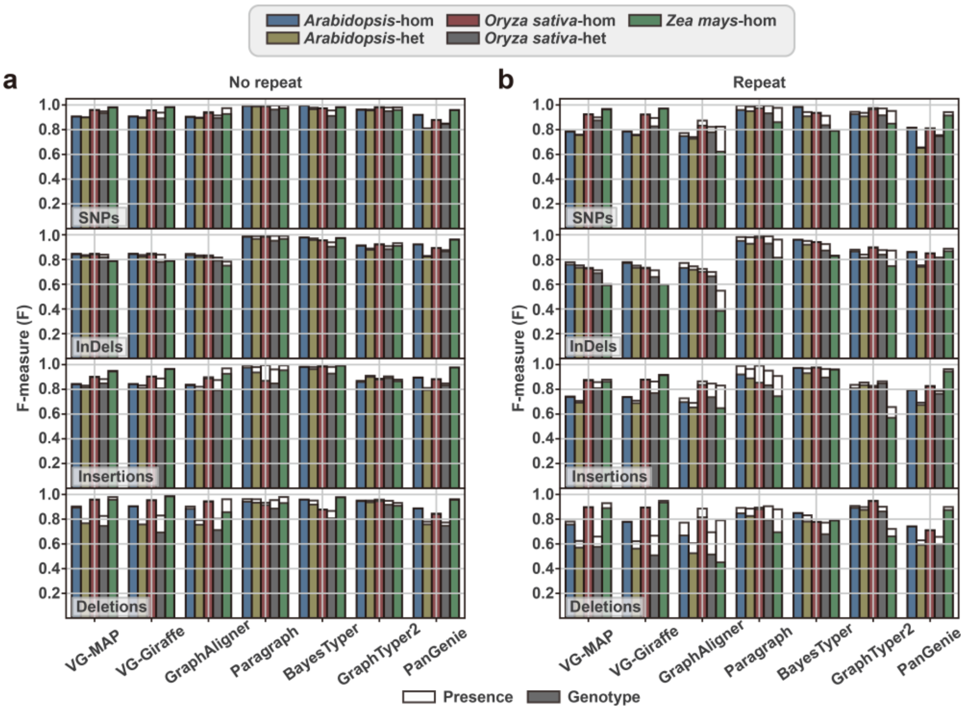


**Figure S5. Effects of genome size and complexity on genotyping.** According to the position of repeats, variants are either from "No repeat" regions **(a)** or from "repeat" regions **(b)**. The genome graph for genotyping is constructed from one reference genome and seven alternative genomes derived by introducing known variants into the reference genome. Both homozygous (hom) and heterozygous (het) alternative genomes of *A. thaliana* and *O. sativa* are simulated. Paired-end (2×150 bp) short reads with 30× depth are simulated for genotyping. Transparent and solid bars represent the ability to predict variant presence and exact genotype.

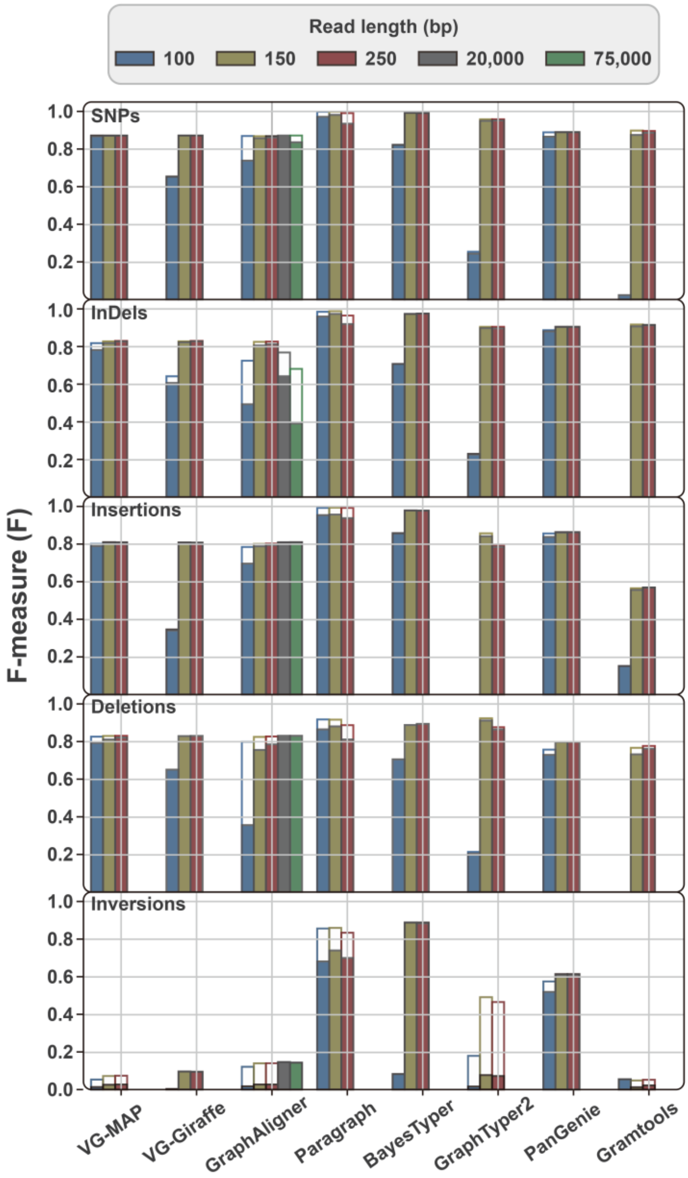


**Figure S6. Impact of read length on genotyping performance based on simulated whole-genome resequencing data of *A. thaliana* genome, partitioned by variant type: SNPs, indels, deletions, insertions, inversions.** The genome graph for genotyping is constructed from one reference genome and seven alternative genomes derived by introducing known variants into the reference genome. Paired-end short-reads with different read lengths and sequencing depth of 30× are simulated for variant genotyping. Two types of long reads are also simulated for genotyping.

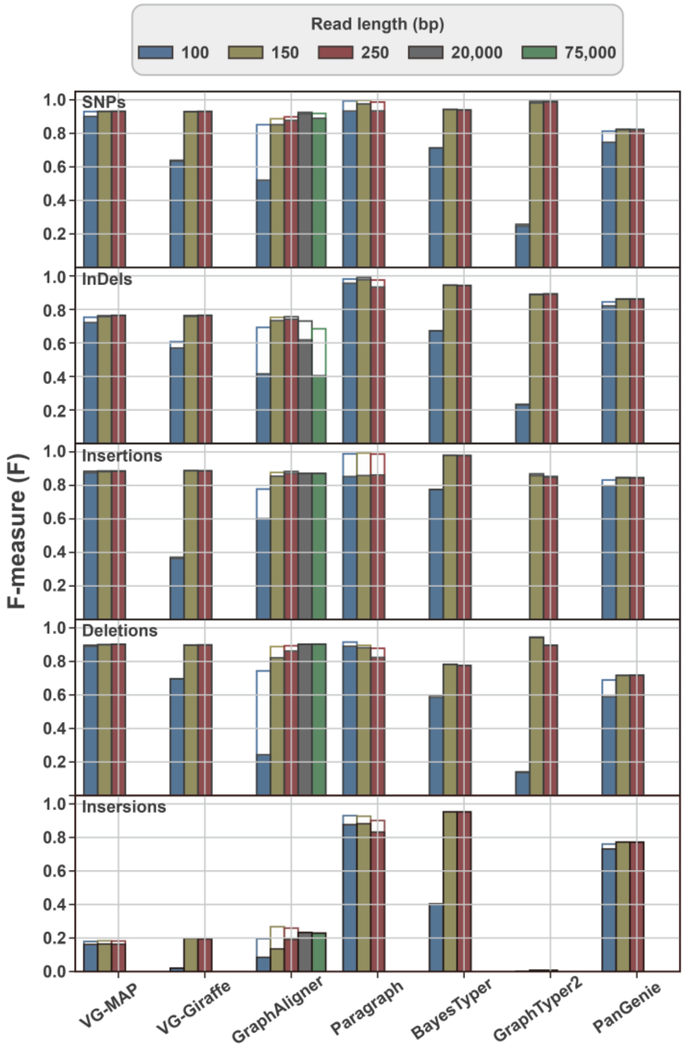


**Figure S7. Impact of read length on genotyping performance based on simulated whole-genome resequencing data of rice genome, partitioned by variant type: SNPs, indels, deletions, insertions, inversions.** The genome graph for genotyping is constructed from one reference genome and seven alternative genomes derived by introducing known variants into the reference genome. Paired-end short-reads with different read lengths and sequencing depth of 30× are simulated for variant genotyping. Two types of long reads are also simulated for genotyping.

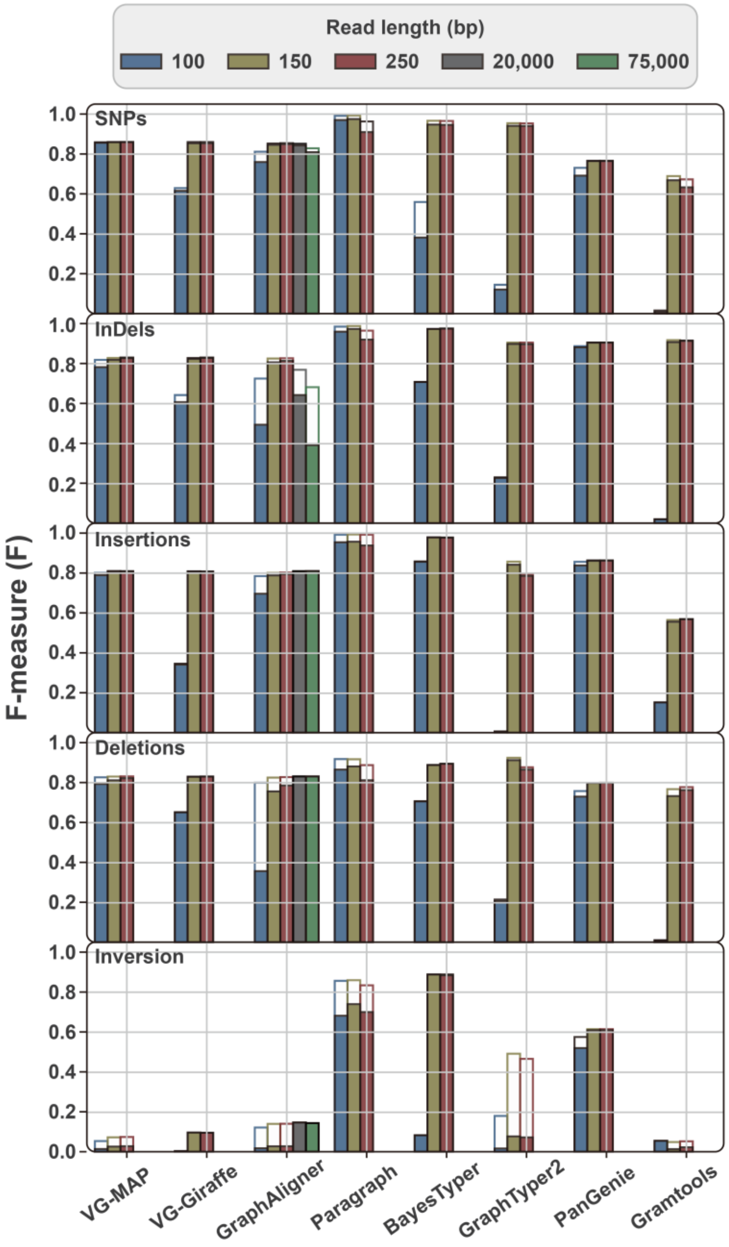


**Figure S8. Impact of read length on genotyping performance based on simulated whole-genome resequencing data of synthetic heterozygous *A. thaliana* genome, partitioned by variant type: SNPs, indels, deletions, insertions, inversions.** The genome graph for genotyping is constructed from one reference genome and seven alternative genomes derived by introducing known variants into the reference genome. Paired-end short-reads with different read lengths and sequencing depth of 30× are simulated for variant genotyping. Two types of long reads are also simulated for genotyping.

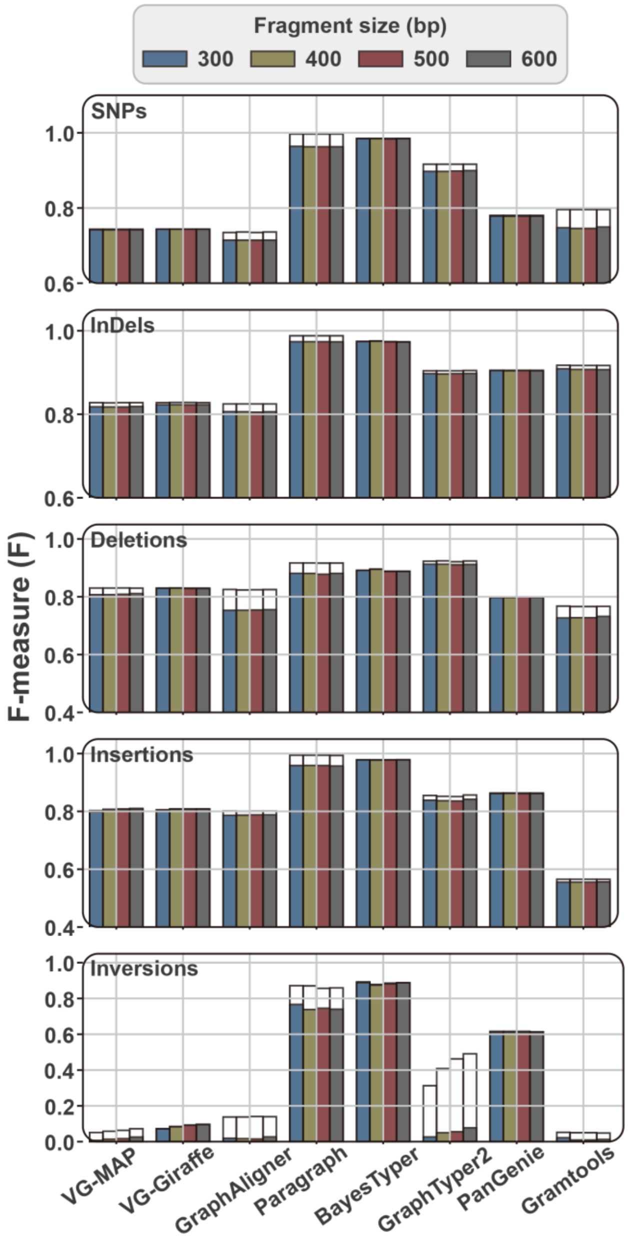


**Figure S9. Impact of fragment size on genotyping performance based on simulated whole-genome resequencing data of synthetic heterozygous *A. thaliana* genome, partitioned by variant type: SNPs, indels, deletions, insertions, inversions.** The genome graph for genotyping is constructed from one reference genome and seven alternative genomes derived by introducing known variants into the reference genome. Paired-end short-reads from DNA fragments with different lengths are simulated for variant genotyping. Paired-end short-reads are simulated with read length of 2×150bp and sequencing depth of 30×.

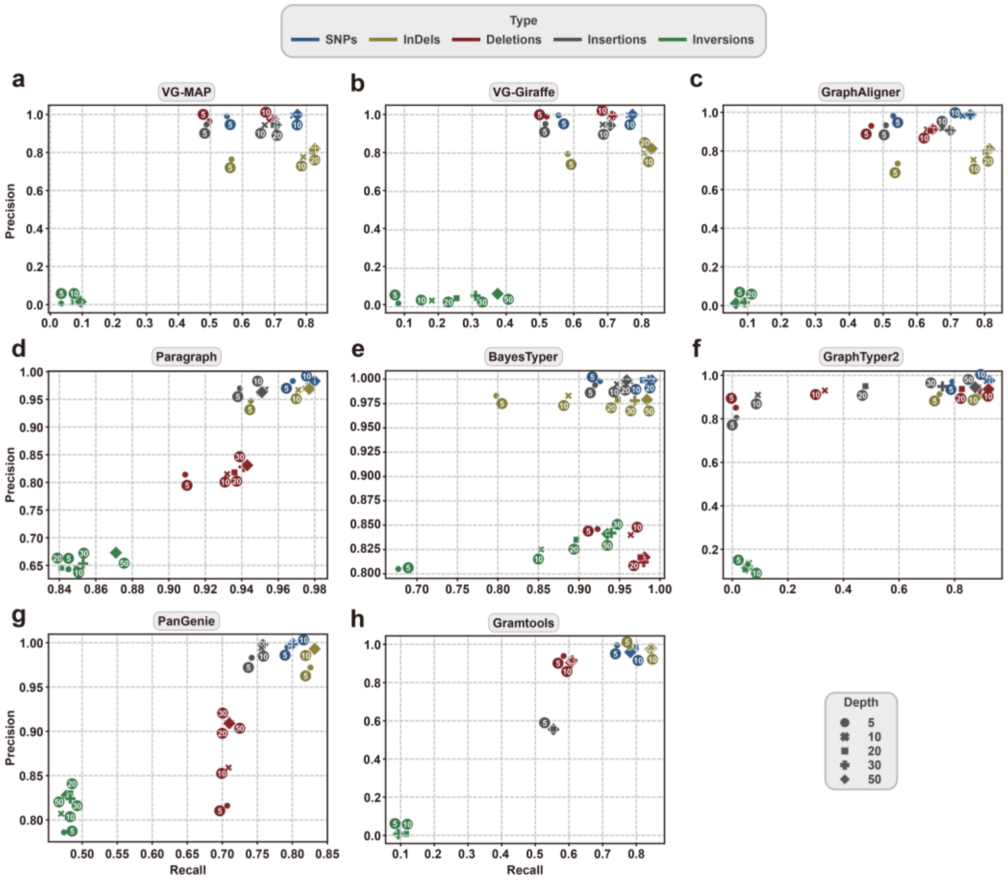


**Figure S10. Performance of variant genotyping under different sequencing depth for eight graph-based genotypers.** The genome graph for genotyping is constructed from the *A. thaliana* reference genome (from accession of Col-0, TAIR10 version) and seven alternative genomes derived by introducing known variants into the reference genome. Variations are represented by different colors: SNPs (blue), indels (yellow), deletions (red), insertions (grey), inversions (green). Paired-end short-reads (read length: 2×150bp) are simulated for variant genotyping.

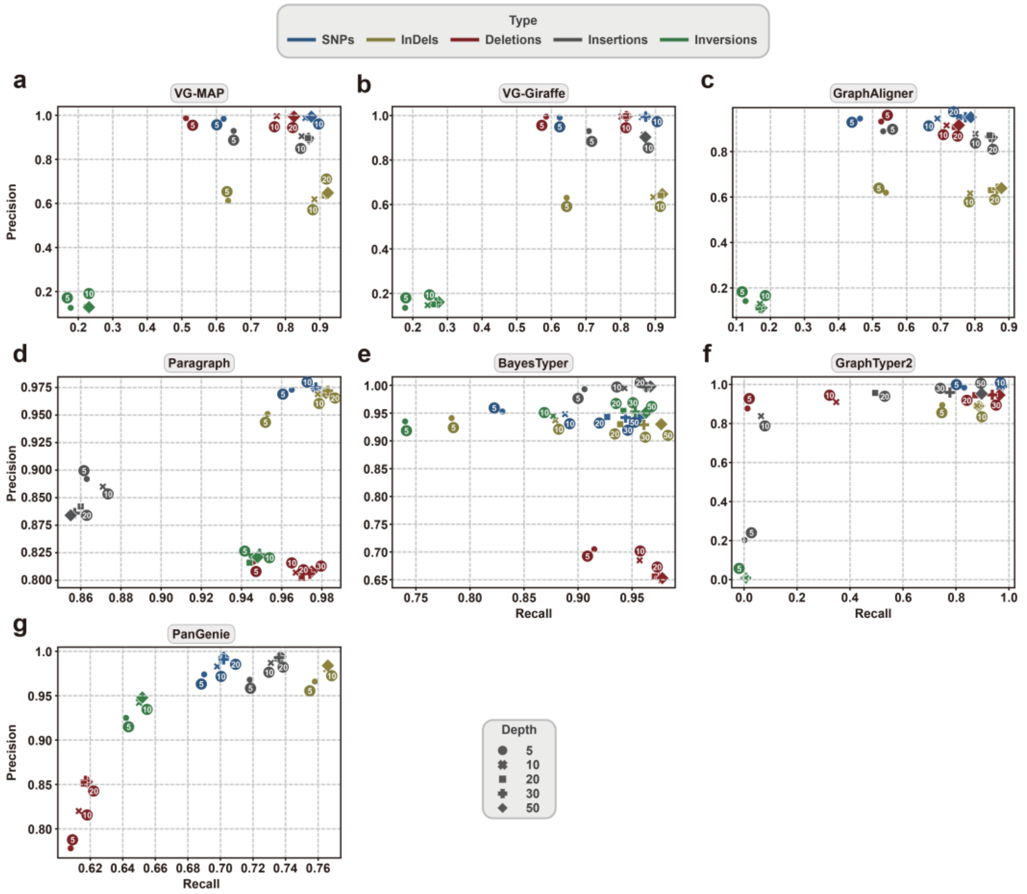


**Figure S11. Performance of variant genotyping under different sequencing depth for eight graph-based genotypers.** The genome graph for genotyping is constructed from the *Oryza sativa* reference genome (from the accession Nipponbare, IGRSP-1.0 version) and seven alternative genomes. Variations are represented by different colors: SNPs (blue), indels (yellow), deletions (red), insertions (grey), inversions (green). Paired-end short-reads (read length: 2×150 bp) are simulated for variant genotyping.

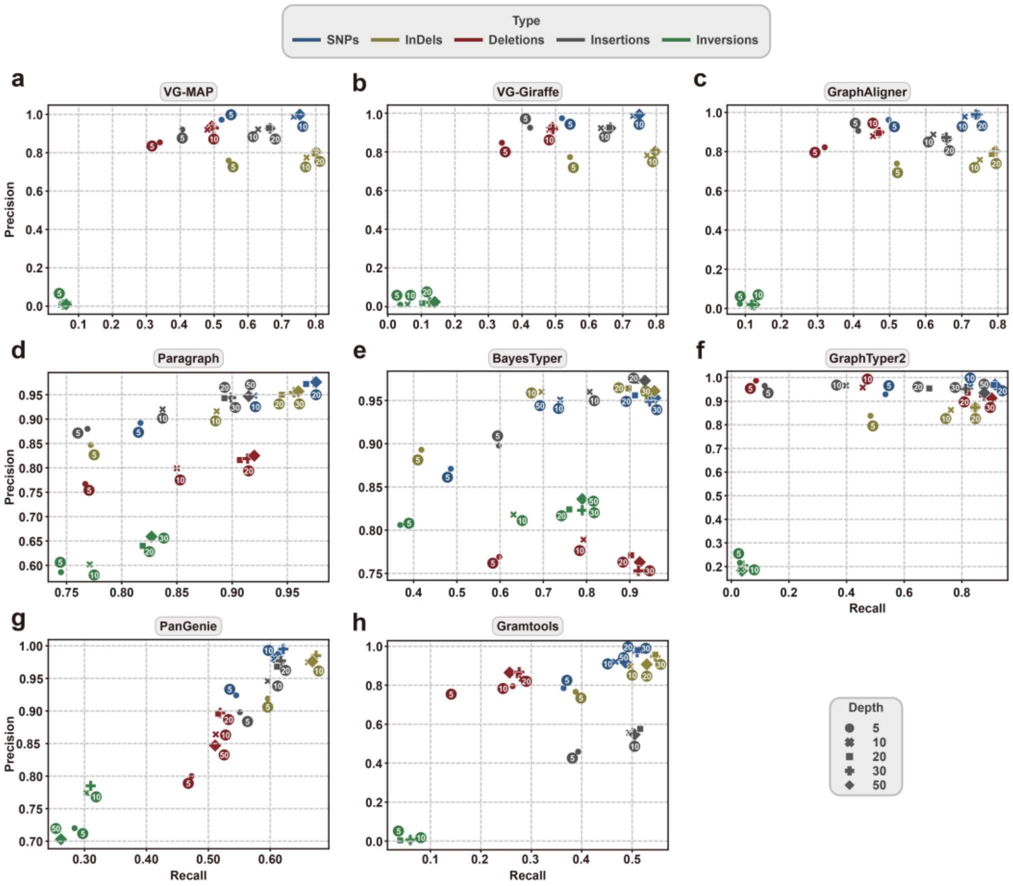


**Figure S12. Performance of variant genotyping under different sequencing depth for eight graph-based genotypers.** The genome graph for genotyping is constructed from the *A. thaliana* reference genome (from the accession of Col-0, TAIR10 version) and seven synthetic heterozygous genomes derived by introducing known variants into the reference genome. Variations are represented by different colors: SNPs (blue), indels (yellow), deletions (red), insertions (grey), inversions (green). Paired-end short-reads (read length: 2×150 bp) are simulated for variant genotyping.

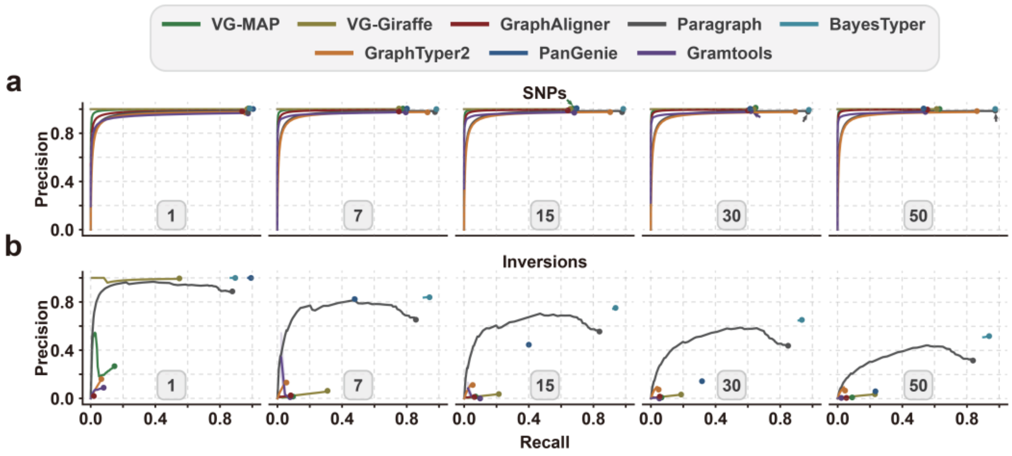


**Figure S13. The effect of genome number on the genotyping performance of different methods, partitioned by variant type: (a) SNPs, (b) inversions.** The five ROC curves correspond to different genome numbers (1, 7, 15, 30, 50). The genome graph for genotyping is constructed from the *A. thaliana* reference genome and different number of alternative genomes derived by introducing known variants into the reference genome. Paired-end short-reads (read length: 2×150bp, sequencing depth: 30×) are simulated for variant genotyping. For each genotyper, precision is plotted against recall as the genotyping quality threshold varies. Read depth (DP) on variant sites is used as a proxy score when genotype quality (GQ) is not available. Arrows indicate the circles hidden by other circles in the plot due to identical or nearly identical precision values.

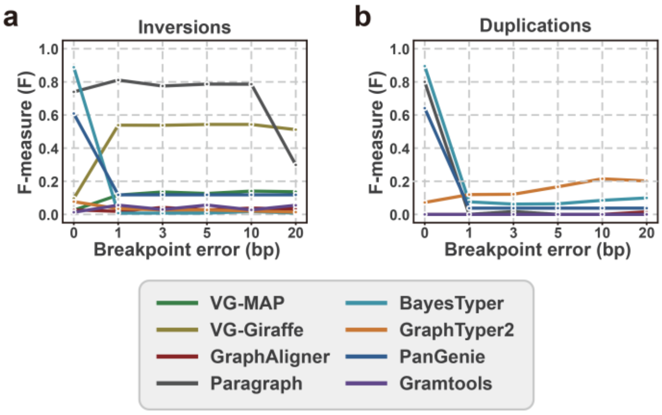


**Figure S14. The impact of breakpoint errors of variation SVs on genotyping performance, partitioned by variant type: (a) inversions and (b) duplications.** The genome graph for genotyping is constructed from the *A. thaliana* reference genome and seven alternative genomes derived by introducing known variants into the reference genome. Paired-end short-reads (read length: 2×150 bp, sequencing depth: 30×) are simulated for variant genotyping.

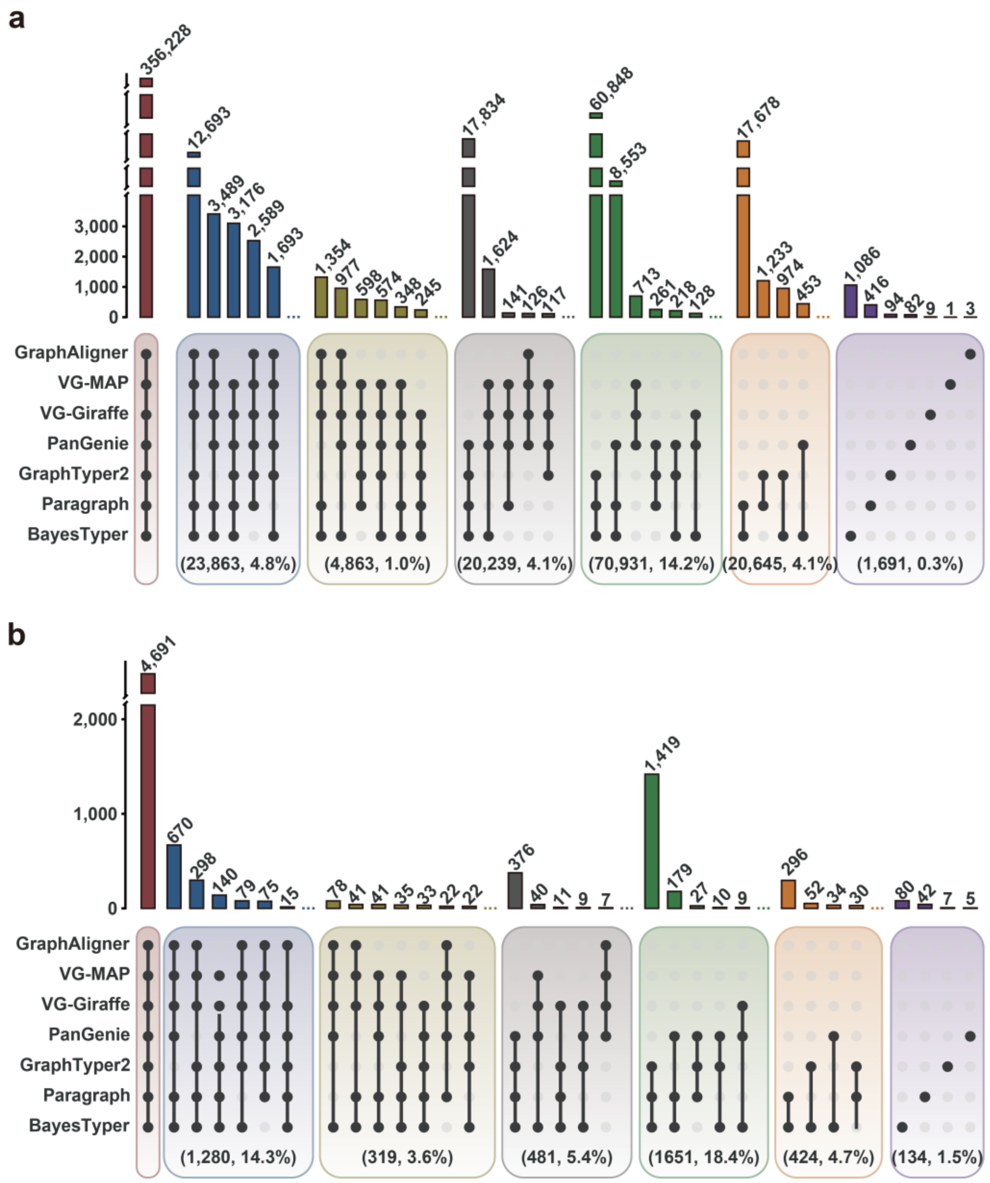


**Figure S15. The intersection of correct genotyping results among different software.** The number of intersections for a specific combination is shown above the column. The number and proportions are provided in parentheses at the bottom. **(a)** The intersection among different software for genotyping SNPs and indels. **(b)** The intersection among different software for genotyping SVs.

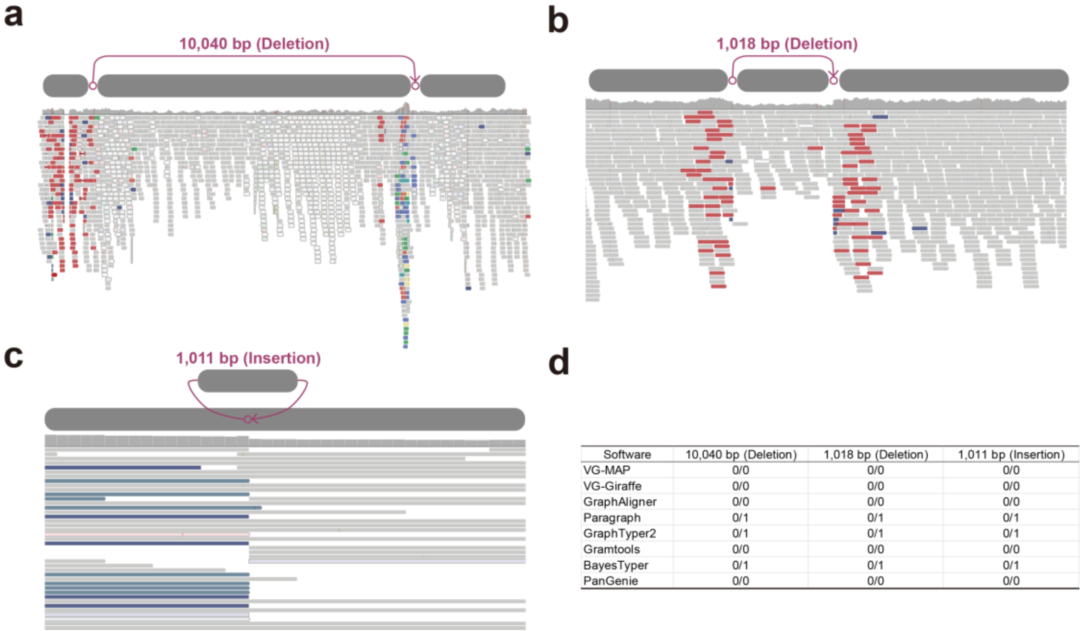


| Software | 10,040 bp (Deletion) | 1,018 bp (Deletion) | 1,011 bp (Insertion) |
|---|---|---|---|
| VG-MAP | 0/0 | 0/0 | 0/0 |
| VG-Giraffe | 0/0 | 0/0 | 0/0 |
| GraphAligner | 0/0 | 0/0 | 0/0 |
| Paragraph | 0/1 | 0/1 | 0/1 |
| GraphTyper2 | 0/1 | 0/1 | 0/1 |
| Gramtools | 0/0 | 0/0 | 0/0 |
| BayesTyper | 0/1 | 0/1 | 0/1 |
| PanGenie | 0/0 | 0/0 | 0/0 |

**Figure S16. Examples of variants correctly genotyped by some tools but not by others.** Two heterozygous (0/1) deletions of **(a)** 10,040 bp and **(b)** 1,018 bp. They are correctly genotyped by Paragraph, GraphTyper2, and BayesTyper. **(c)** A heterozygous (0/1) insertion with a length of 1,011 bp. **(d)** Genotyping result of these three variants.

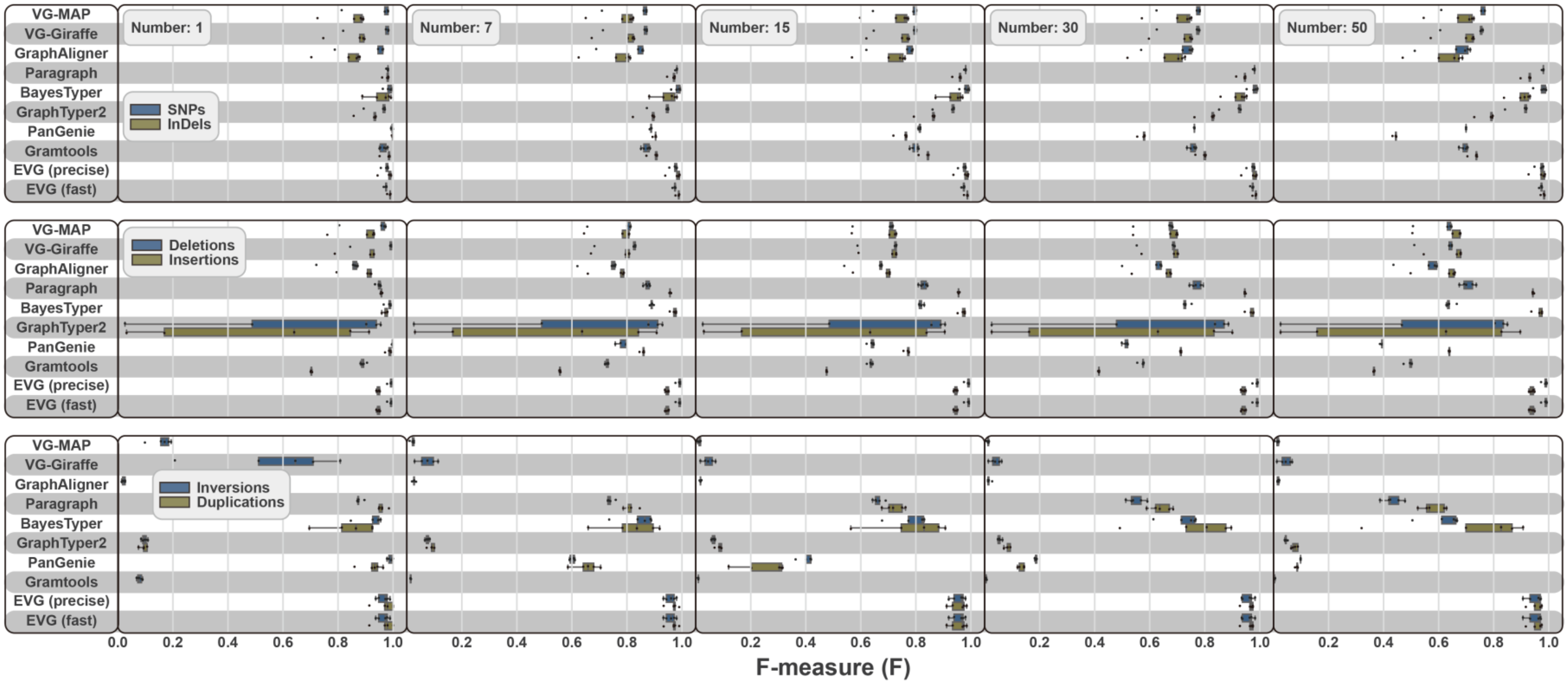


**Figure S17. The influence of graphed genome number and sequencing depth on the genotyping performance of EVG, partitioned by variant type: SNPs, indels, deletions, insertions, inversions, duplications.** The genome graph for genotyping is constructed from the *A. thaliana* reference genome and seven alternative genomes derived by introducing known variants into the reference genome. Paired-end short-reads with read length 2×150 bp are simulated for variant genotyping. The five points in each bar chart correspond to different sequencing depth (5×, 10×, 20×, 30×, 50×).

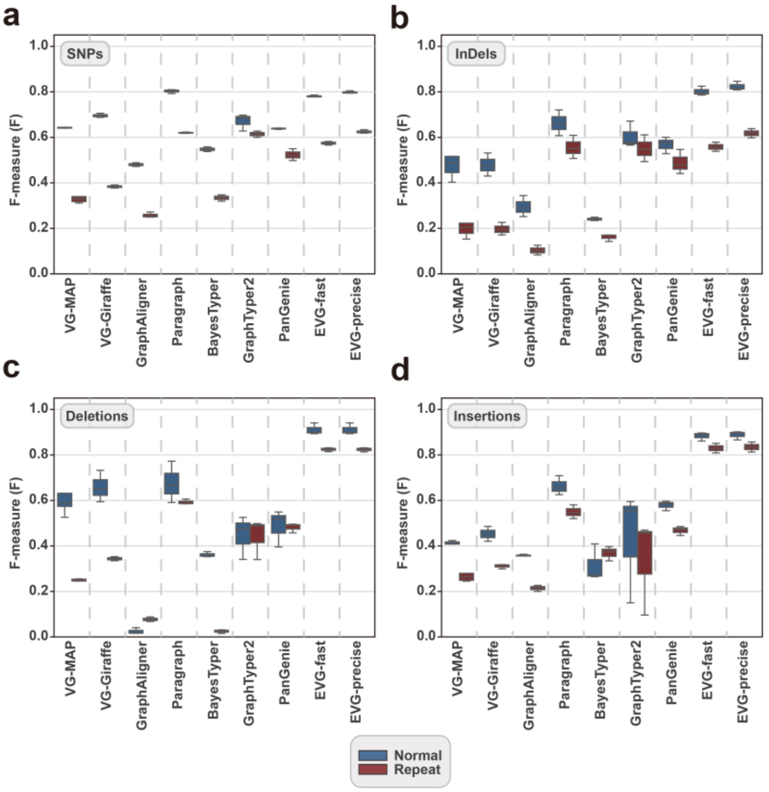


**Figure S18 Effects of repetitive sequences on variant genotyping in maize, partitioned by variant type: SNPs, indels, deletions, insertions.** The genome graph for genotyping is constructed from the maize reference genome and seven alternative genomes derived by introducing known variants into the reference genome. Paired-end short-reads with read length of 2×150 bp and sequencing depth of 30× are simulated for variant genotyping.

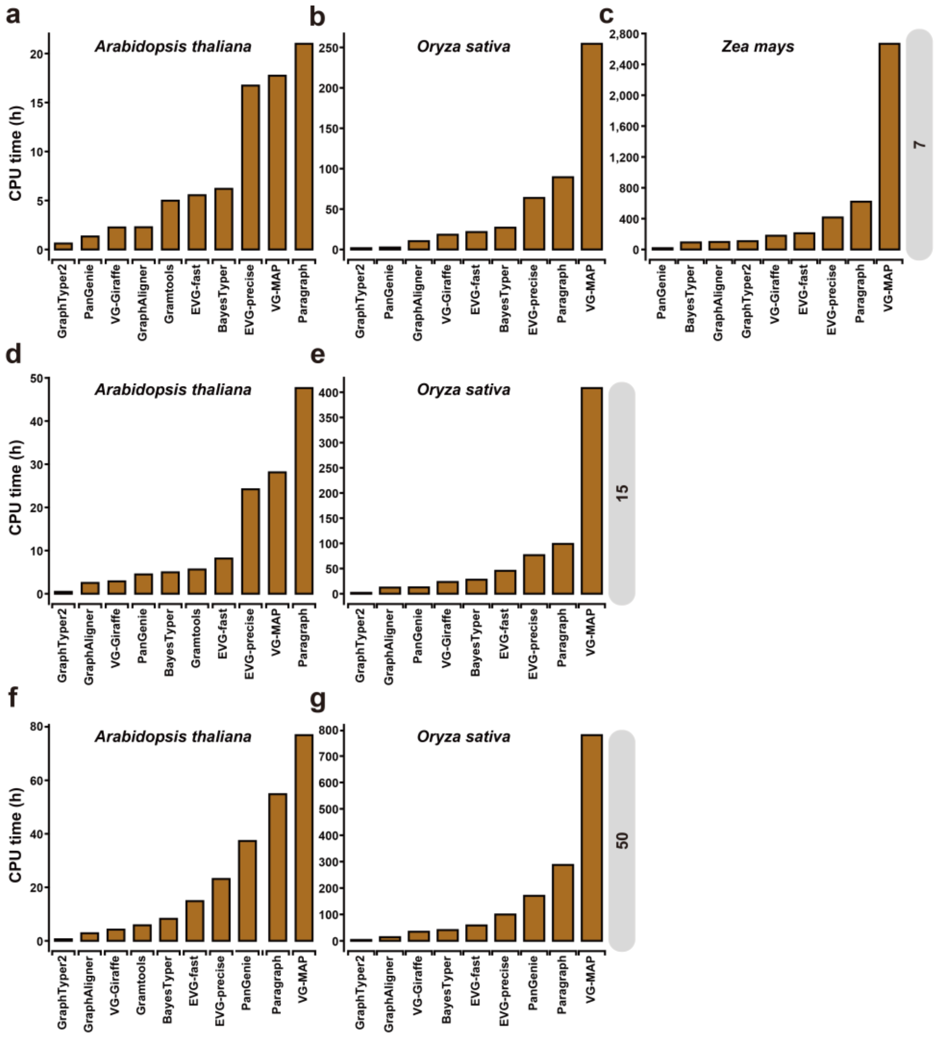


**Figure S19. Runtime usage for variant genotyping in different plant genomes.** Total runtime is measured under using 10 threads. The genome graph for genotyping is constructed from one reference genome and different number (7 for (a, b, c), 15 for (d, e), 50 for (f, g)) of alternative genomes derived by introducing known variants into the reference genome. Paired-end short-reads with read length of 2×150 bp and sequencing depth of 30× are simulated for variant genotyping.

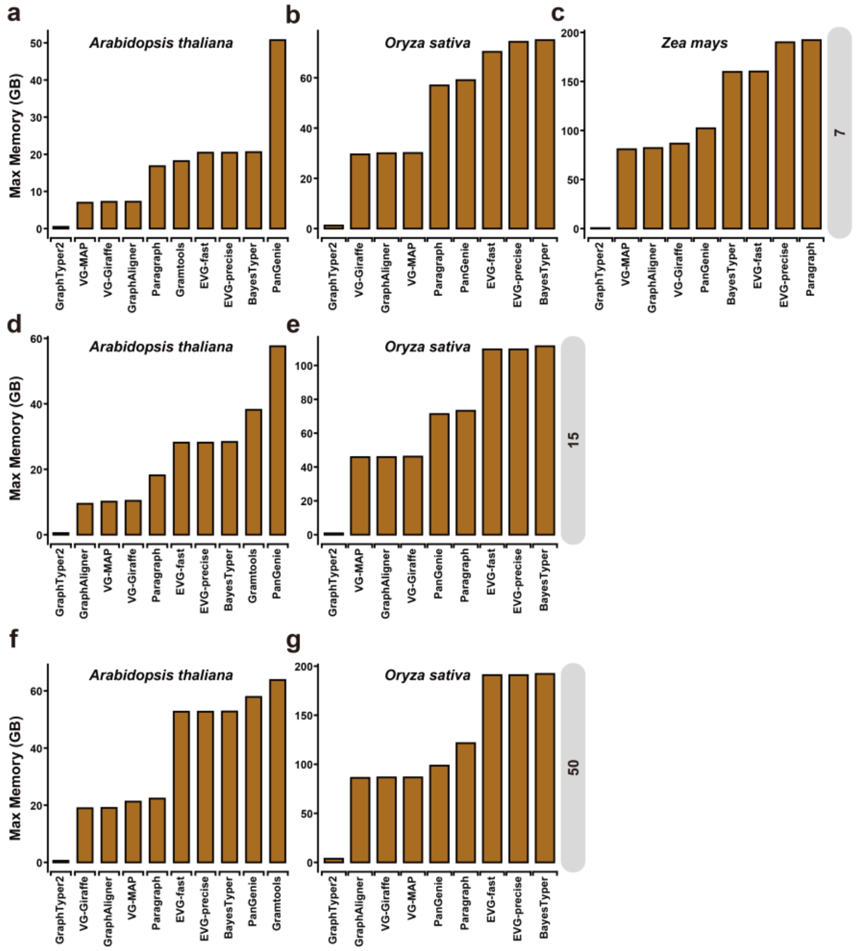


**Figure S20. Memory usage.** Peak memory usage is measured under using 10 threads. The genome graph for genotyping is constructed from one reference genome and different numbers (7 for (a, b, c), 15 for (d, e), 50 for (f, g)) of alternative genomes derived by introducing known variants into the reference genome. Paired-end short-reads with read length of 2×150 bp and sequencing depth of 30× are simulated for variant genotyping.

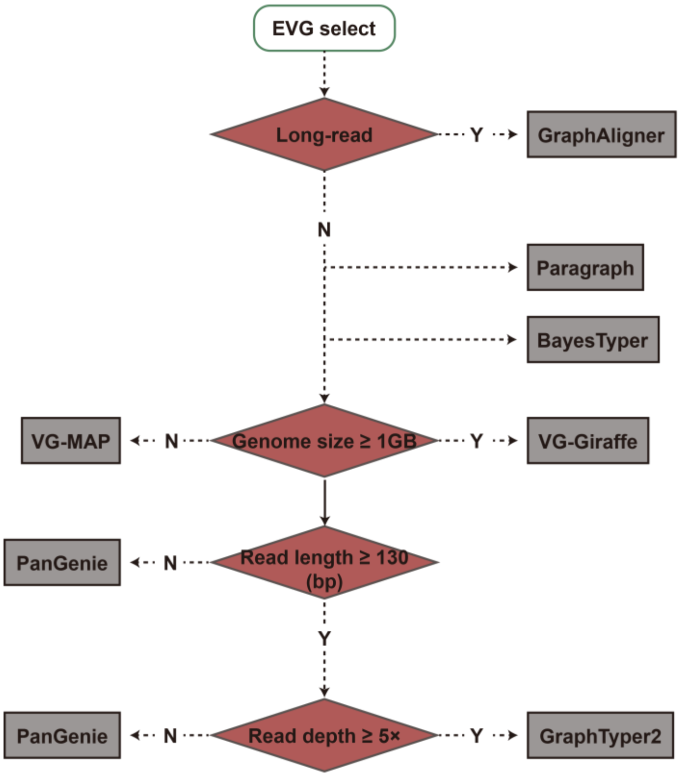


**Figure S21. The process of software selection in the EVG pipeline.**

**Table S1. Overview of graph genome tools.**

| Software | Graph types | Indexing | Mapping | Genotyping | Reference |
|---|---|---|---|---|---|
| VG-MAP | Variation graph (bidirected) | GCSA2 | Read alignment (Seed-cluster-chain, GSSW) | Coverage of local haplotypes | [1] |
| BayesTyper | Variation graph (DAG) | k-mer hash table | Read k-mer (n-best) | Generative model of the sequencing process with noise and diplotype k-mer counts | [2] |
| Paragraph | Variation graph (DAG) | k-mer hash table | Read alignment (S&E, GSSW) | Relative likelihood of local haplotypes based on coverages of breakpoints | [3] |
| GraphTyper2 | Variation graph (DAG) | k-mer hash table | Read alignment (S&E) | Relative likelihood of local haplotypes based on breakpoint coverages and coverage in/decrease | [4, 5] |
| VG-Giraffe | Variation graph (bidirected) | GBWT, Minimizer | Read alignment (Seed-cluster-chain, GBWT) | Coverage of local haplotype | [6] |
| Gramtools | Variation graph (nested DAG) | vBWT | Read alignment (S&E) | Likelihood of allele based on base-level and allele-level coverage | [7] |
| PanGenie | Variation graph (DAG) | k-mer hash table | Read k-mer | HMM of local haplotypes with k-mer counts and recombination rates | [8] |
| GraphAligner (+ VG) | Variation or overlap graph (bidirected) | Minimizer (default) | Read alignment (S&E) | VG used | [9] |
| HISAT-genotype | Variation graph (DAG) | hierarchical GFM (HGFM), FM index, Minimizer | Read alignment (S&E) | EM algorithm | [10] |
| Minos | Variation graph (nested DAG) | vBWT | Read alignment (S&E) | Gramtools used | [11] |
| KAGE | Variation graph (DAG) | k-mer hash table | Read k-mer | Poisson distribution | [12] |

| | | | | | |
|---|---|---|---|---|---|
| Seven Bridges Genomics | Variation graph (DAG) | k-mer hash table | Read alignment (S&E, SIMD) | HMM | [13] |

Note: DAG: directed acyclic graph, S&E: Seed-and-Extend, GCSA: Generalized Compressed Suffix Array, GBWT: Graph Burrows-Wheeler Transform, HMM: Hidden Markov Model, vBWT: variation- aware Burrows-Wheeler Transform.

**Table S2. Summary of reference genomes used in this study.**

| Organism | Karyotype | Accession | Genome size (Mb) | Repeat Content (%) |
|---|---|---|---|---|
| *Arabidopsis thaliana* | 2n = 2× = 10 | Col-0 | 135.0 | 21.4 |
| *Oryza sativa* | 2n = 2× = 24 | Nipponbare | 410.0 | 58.0 |
| *Zea mays* | 2n = 2× = 20 | B73 | 2300.0 | 88.9 |
| *Brassica napus* | 2n = 4× = 38 | ZS11 | 1200.0 | 65.6 |
| *Prunus armeniaca* | 2n = 2× = 16 | Rojo Pasión | 243.9 | 45.0 |

**Table S3. Summary of simulated short reads for variant genotyping.**

| Organism | Heterozygosity rate (%) | Read length (bp) | Fragment size (bp) | Depth (×) | Read number (bp) | Read base (bp) |
|---|---|---|---|---|---|---|
| *Arabidopsis thaliana* | 0 | 2×100 | 400 | 30 | 36,012,826 | 3,601,282,600 |
| *Arabidopsis thaliana* | 0 | 2×150 | 300 | 30 | 24,008,504 | 3,601,275,600 |
| *Arabidopsis thaliana* | 0 | 2×150 | 400 | 30 | 24,008,466 | 3,601,269,900 |
| *Arabidopsis thaliana* | 0 | 2×150 | 500 | 30 | 24,008,436 | 3,601,265,400 |
| *Arabidopsis thaliana* | 0 | 2×150 | 600 | 5 | 4,001,376 | 600,206,400 |
| *Arabidopsis thaliana* | 0 | 2×150 | 600 | 10 | 8,002,792 | 1,200,418,800 |
| *Arabidopsis thaliana* | 0 | 2×150 | 600 | 20 | 16,005,588 | 2,400,838,200 |
| *Arabidopsis thaliana* | 0 | 2×150 | 600 | 30 | 24,008,372 | 3,601,255,800 |
| *Arabidopsis thaliana* | 0 | 2×150 | 600 | 50 | 40,014,040 | 6,002,106,000 |
| *Arabidopsis thaliana* | 0 | 2×250 | 600 | 30 | 14,404,844 | 3,601,211,000 |
| *Arabidopsis thaliana* | 0.27 | 2×150 | 600 | 30 | 23,938,416 | 3,590,762,400 |
| *Arabidopsis thaliana* | 0.52 | 2×150 | 600 | 30 | 23,950,450 | 3,592,567,500 |
| *Arabidopsis thaliana* | 1.03 | 2×150 | 600 | 30 | 23,913,474 | 3,587,021,100 |
| *Arabidopsis thaliana* | 2.07 | 2×150 | 600 | 30 | 23,902,250 | 3,585,337,500 |
| *Arabidopsis thaliana* | 2.35 | 2×150 | 600 | 30 | 23,894,154 | 3,584,123,100 |
| *Oryza sativa* | 0 | 2×100 | 400 | 30 | 106,813,926 | 10,681,392,600 |
| *Oryza sativa* | 0 | 2×150 | 600 | 5 | 11,866,870 | 1,780,030,500 |
| *Oryza sativa* | 0 | 2×150 | 600 | 10 | 23,733,388 | 3,560,008,200 |
| *Oryza sativa* | 0 | 2×150 | 600 | 20 | 47,467,192 | 7,120,078,800 |
| *Oryza sativa* | 0 | 2×150 | 600 | 30 | 71,200,086 | 10,680,012,900 |
| *Oryza sativa* | 0 | 2×150 | 600 | 50 | 118,667,134 | 17,800,070,100 |
| *Oryza sativa* | 0 | 2×250 | 600 | 30 | 42,714,776 | 10,678,694,000 |
| *Oryza sativa* | 0.34 | 2×150 | 600 | 30 | 72,548,332 | 10,882,249,800 |
| *Zea mays* | 0 | 2×150 | 600 | 30 | 432,049,692 | 64,807,453,800 |
| *Brassica napus* | 0 | 2×150 | 600 | 30 | 191,710,668 | 28,756,600,200 |

**Table S4. The number of genomes and variants included in different genome graphs based on the simulated datasets.**

| Organism | Heterozygosity Rate (%) | Genome Number | SNPs | Indels | SVs |
|---|---|---|---|---|---|
| *Arabidopsis thaliana* | 0 | 1 | 467,512 | 38,207 | 9,036 |
| *Arabidopsis thaliana* | 0 | 7 | 1,202,564 | 100,000 | 18,369 |
| *Arabidopsis thaliana* | 0 | 15 | 1,797,784 | 176,257 | 28,562 |
| *Arabidopsis thaliana* | 0 | 30 | 2,313,019 | 266,732 | 44,174 |
| *Arabidopsis thaliana* | 0 | 50 | 2,942,438 | 364,524 | 67,726 |
| *Arabidopsis thaliana* | 0.27 | 7 | 821,040 | 78,150 | 19,195 |
| *Arabidopsis thaliana* | 0.52 | 7 | 1,211,898 | 113,220 | 19,761 |
| *Arabidopsis thaliana* | 1.03 | 7 | 1,638,958 | 253,322 | 19,782 |
| *Arabidopsis thaliana* | 2.07 | 7 | 2,979,610 | 259,044 | 19,740 |
| *Arabidopsis thaliana* | 2.35 | 7 | 3,238,944 | 466,287 | 19,637 |
| *Oryza sativa* | 0 | 1 | 1,348,007 | 58,898 | 13,745 |
| *Oryza sativa* | 0 | 7 | 1,764,132 | 104,635 | 34,729 |
| *Oryza sativa* | 0 | 15 | 2,251,267 | 160,311 | 64,260 |
| *Oryza sativa* | 0 | 30 | 3,007,606 | 246,960 | 110,511 |
| *Oryza sativa* | 0 | 50 | 3,727,286 | 331,631 | 156,519 |
| *Oryza sativa* | 0.34 | 7 | 1,944,334 | 111,037 | 38,074 |
| *Zea mays* | 0 | 7 | 5,766,579 | 120,687 | 126,353 |
| *Brassica napus* | 0 | 7 | 1,055,562 | 122,970 | 153,796 |

**Table S5. The number of alternative genomes and variants included in different genome graphs based on the real dataset.**

| Organism | Sample | Genome Number | SNPs | Indels | SVs | Reference |
|---|---|---|---|---|---|---|
| *Arabidopsis thaliana* | An-1 | 1 | 686,989 | 51,369 | 2,245 | [14] |
| *Arabidopsis thaliana* | C24 | 1 | 793,416 | 56,863 | 2,372 | [14] |
| *Arabidopsis thaliana* | Cvi-0 | 1 | 931,980 | 66,797 | 2,891 | [14] |
| *Arabidopsis thaliana* | An-1, C24, Cvi-0, Eri, Kyo, Ler, Sha | 7 | 2,512,655 | 219,797 | 11,816 | [14] |
| *Oryza sativa* | TG19 | 1 | 481,625 | 41,347 | 2,935 | [15] |
| *Oryza sativa* | TG28 | 1 | 3,067,554 | 218,206 | 9,058 | [15] |
| *Oryza sativa* | TG78 | 1 | 3,097,123 | 225,438 | 8,708 | [15] |
| *Oryza sativa* | TG19, TG28, TG78, TG81, TG63, TG80, TG8 | 7 | 4,264,744 | 347,690 | 35,877 | [15] |
| *Zea mays* (chr10) | B97 | 1 | 787,042 | 19,882 | 1,717 | [16, 17] |
| *Zea mays* (chr10) | CML52 | 1 | 795,612 | 25,558 | 1,911 | [16, 17] |
| *Zea mays* (chr10) | CML69 | 1 | 902,189 | 30,600 | 1,797 | [16, 17] |
| *Zea mays* (chr10) | B97, CML52, CML69, CML103, CML228, CML247, CML277 | 7 | 1,940,430 | 84,894 | 8,596 | [16, 17] |
| *Prunus armeniaca* | Rojo Pasión | 1 | 1,328,299 | 44,796 | 5,997 | [18] |
| *Prunus armeniaca* | Rojo Pasión, A02, A04, B03, C04, E02, H18 | 7 | 5,617,665 | 146,865 | 6,356 | [18, 19] |

**Table S6. Summary of real short read datasets for variant genotyping.**

| Organism | Line | Depth (×) | Read number (bp) | Read length (bp) | Read base (bp) | Reference |
|---|---|---|---|---|---|---|
| *Arabidopsis thaliana* | An-1 | 30 | 35,582,684 | 100 | 3,574,388,282 | [14] |
| *Arabidopsis thaliana* | C24 | 30 | 35,550,956 | 100 | 3,574,382,394 | [14] |
| *Arabidopsis thaliana* | Cvi-0 | 30 | 35,583,672 | 100 | 3,574,357,424 | [14] |
| *Oryza sativa* | TG19 | 30 | 75,039,376 | 149 | 11,196,786,703 | [15] |
| *Oryza sativa* | TG28 | 30 | 75,159,944 | 148 | 11,197,441,298 | [15] |
| *Oryza sativa* | TG79 | 30 | 75,138,760 | 149 | 11,197,018,772 | [15] |
| *Zea mays* | B97 | 30 | 437,373,216 | 146 | 63,955,692,185 | [16] |
| *Zea mays* | CML52 | 30 | 402,746,050 | 148 | 59,629,814,562 | [16] |
| *Zea mays* | CML69 | 30 | 439,206,688 | 145 | 63,958,356,137 | [16] |
| *Prunus armeniaca* | Rojo Pasión | 30 | 42,715,012 | 150 | 6,407,251,800 | [18] |
